# CONSTITUTIVE OPENING OF THE Kv7.2 PORE ACTIVATION GATE CAUSES *KCNQ2*-DEVELOPMENTAL ENCEPHALOPATHY

**DOI:** 10.1101/2024.05.20.593680

**Authors:** Mario Nappi, Giulio Alberini, Alessandro Berselli, Agnese Roscioni, Maria Virginia Soldovieri, Vincenzo Barrese, Sarah Weckhuysen, Ting-Gee Annie Chiu, Ingrid E. Scheffer, Fabio Benfenati, Luca Maragliano, Francesco Miceli, Maurizio Taglialatela

**Affiliations:** Dept. of Neuroscience, University of Naples “Federico II”, 80131 Naples, IT; Center for Synaptic Neuroscience and Technology, Istituto Italiano di Tecnologia, 16132 Genova, IT; IRCCS Ospedale Policlinico San Martino, 16132 Genova, IT; Department of Experimental Medicine, Università degli Studi di Genova, 16132 Genova, IT; Department of Life and Environmental Sciences, Polytechnic University of Marche, 60131 Ancona, IT; Dept. of Medicine and Health Science, University of Molise, 86100 Campobasso, IT; Applied & Translational Neurogenomics Group, Vlaams Instituut voor Biotechnology (VIB) Center for Molecular Neurology, VIB, Antwerp 2610, BE; Translational Neurosciences, Faculty of Medicine and Health Science, University of Antwerp, Antwerp 2610, BE; Department of Neurology, Antwerp University Hospital, Antwerp 2610, BE; µNEURO Research Centre of Excellence, University of Antwerp, Antwerp 2610, BE; University of Melbourne, Austin Health, Melbourne, AU; University of Melbourne, Austin and Royal Children’s Hospital, Florey and Murdoch Children’s Research Institutes, Melbourne, AU; These authors contributed equally: Mario Nappi, Giulio Alberini

**Keywords:** Potassium channels, developmental and epileptic encephalopathies, genotype-phenotype correlations, *KCNQ*, gene variants

## Abstract

Pathogenic variants in *KCNQ2* encoding for Kv7.2 voltage-gated potassium channel subunits cause developmental encephalopathies (*KCNQ2*-encephalopathies), both with and without epilepsy. We herein describe the clinical, *in vitro* and *in silico* features of two encephalopathy-causing variants (A317T, L318V) in Kv7.2 affecting two consecutive residues in the S_6_ activation gate undergoing large structural rearrangements during pore opening. Currents through these mutant channels displayed increased density, hyperpolarizing shifts in activation gating, and insensitivity to phosphatidylinositol 4,5-bisphosphate (PIP_2_), a critical regulator of Kv7 channel function; all these features are consistent with a strong gain-of-function effect. An increase in single-channel open probability, with no change in membrane abundance or single-channel conductance, was responsible for the observed gain-of-function effects. All-atoms Molecular Dynamics simulations revealed that the mutations widened the inner pore gate and stabilized a constitutively open channel configuration in the closed state, with minimal effects on the open conformation. Thus, a PIP_2_-independent stabilization of the inner pore gate open configuration is a novel molecular pathogenetic mechanism for *KCNQ2*-developmental encephalopathies.

## INTRODUCTION

Variants in *KCNQ2* or *KCNQ3* genes, encoding for Kv7.2 and Kv7.3 voltage-gated potassium (K^+^) channel subunits, respectively, can cause a spectrum of mostly neonatal-onset phenotypes ranging from self-limited familial neonatal epilepsy (SLFNE) to developmental and epileptic encephalopathies (DEEs). ^1–5^ DEE are the most severe group of epilepsies, characterized by early-onset drug-resistant seizures, epileptiform activity on EEG, developmental delay or regression, and an increased risk of early death^6^. DEEs often have a genetic etiology identified in 50% of cases, with *KCNQ2* and *KCNQ3* pathogenic variants being responsible for the largest fraction of neonatal-onset DEEs.^7^ More recently, the phenotypic spectrum associated to KCNQ2 variants has further expanded to include intellectual disability (ID) without seizures.^8^

Kv7.2 and Kv7.3 subunits are mainly expressed in neurons where they can assemble as homo- or hetero-tetrameric channels underlying the M-current (I_KM_), a non-inactivating K^+^ current with slow activation and deactivation kinetics, that regulates the resting membrane potential and suppresses repetitive firing ^9,10^. Kv7 subunits contain six transmembrane (TM) segments (S_1_-S_6_), with S_1_-S_4_ forming the voltage-sensing domain (VSD), and S_5_, S_6_ together with the S_5_-S_6_ intervening linker forming the pore domain (PD).^10,11^ This latter can be divided in three structurally- and functionally-distinct regions: two constricted areas located toward the extracellular or the intracellular sides, represented by the selectivity filter (SF) and the activation gate (AG), respectively, and a central cavity (CC) in between.^12^ The cytosolic C-terminal region contains four α-helices (HA, HB, HC and HD) where critical sites for assembly, interaction with regulatory molecules, subcellular localization, and binding of accessory proteins such as calmodulin (CaM)^10^ have been identified. In addition to membrane depolarization, Kv7 channel opening requires endogenous ligands such as the membrane lipid phosphatidylinositol 4,5-bisphosphate (PIP_2_), which binds to distinct channel regions in the VSD and the PD,^13^ and induces large and complex structural rearrangements.^14,15^

Over the years, functional *in vitro* studies in heterologous expression systems revealed that the majority of pathogenic variants in *KCNQ2* or *KCNQ3* cause loss-of-function (LoF) effects by reducing maximal current density, decreasing subunit expression and/or trafficking to the membrane, shifting the activation gating in the depolarizing direction, and/or altering the response to regulatory proteins such as kinases.^2^ A decreased I_KM_ density has been also observed in neurons differentiated from human induced pluripotent stem cells carrying distinct LoF variants associated with *KCNQ2*-DEE.^16^ A correlation between the extent of LoF *in vitro* and the clinical phenotype has been reported, with SLFNE-causing variants being associated with a milder degree of current impairment (haploinsufficiency), whereas most DEE-causing pathogenic variants exert more dramatic, dominant-negative functional consequences.^17,18^ This genotype/phenotype correlation was also observed in neuronal populations from murine models of SFLNE, which exhibit a relatively small decrease in I_KM_ density (∼25%), as compared to a mouse model carrying a Kv7.2 recurrent pathogenic variant causing DEE, in which a ∼50% decrease was found.^19^

In addition to LoF, few heterozygous missense variants enhancing channel function (gain-of-function, GoF) have been described, in both *KCNQ2*^3^ and *KCNQ3*.^4^ For *KCNQ2*, phenotypes associated with GoF variants differ from the classical *KCNQ2*-DEE phenotype, ranging from severe ID without neonatal seizures, but with a characteristic non-epileptic myoclonus and poor prognosis, to moderate/severe ID and infantile epilepsy.^20–23^ Similarly, pathogenic GoF variants in *KCNQ3* are associated with global developmental delay evolving to ID, sleep-activated near-continuous multifocal spikes, and autistic features, without neonatal seizures.^23^ The mechanisms by which *KCNQ2* GoF variants cause such diverse clinical phenotypes are complex and not completely understood. ^2,25^ Differential effects on excitability of distinct neuronal populations of cortical neurons have been recently described in transgenic mice carrying a Kv7.2 GoF variant, with layer 2/3 pyramidal neurons becoming hyperexcitable, and CA1 pyramidal neurons hypoexcitable.^26^

All pathogenic GoF variants are missense and, with one exception (G239S) which affects the PD of Kv7.2,^21^ are located within the VSD of either Kv7.2^20,27,28^ or Kv7.3^24^ subunits. These GoF variants cause a hyperpolarizing shift in voltage-dependent activation gating by stabilizing the activated configuration of S_4_ and impeding VSD repositioning upon hyperpolarization in the physiological voltage range.^29,30^

A correlation between genotype, clinical phenotype and *in vitro* functional consequences has been revealed not only for *KCNQ2*- and *KCNQ3*-related disorders but also for several other channelopathies causing neurodevelopmental disorders;^31,32^ therefore, detailing the specific molecular mechanism(s) by which each variant affects channel function may provide critical insights into phenotypic heterogeneity, prognosis, and may direct precision medicine treatments.

Here, by a translational multidisciplinary approach linking clinical phenotyping to mutagenesis, electrophysiology, biochemistry, and all-atom Molecular Dynamics (MD), we investigated the functional and structural consequences of three missense variants affecting residues in the pore AG of Kv7.2 and Kv7.3 subunits causing severe developmental encephalopathy. These include: 1. the *KCNQ2* A317T variant newly identified in a patient with ID, autism spectrum disorders (ASD), but without seizures; 2. the *KCNQ2* L318V variant found in a DEE-affected patient with infantile epileptic spasms without preceding neonatal seizures^33^, and 3. the *KCNQ3* A356T variant, paralogous to the A317T variant in *KCNQ2*, identified in a large cohort of patients with ID.^34^ Altogether, we found that these variants exert strong GoF effects by stabilizing the open state configuration of the pore domain, thus revealing a new molecular pathogenetic mechanism for *KCNQ2*- and *KCNQ3*-related neurodevelopmental disorders.

## RESULTS

### Clinical and genetic features of the 3 patients included in the present study

*Patient 1* is a 5 year old boy with severe ASD and severe ID. He had global developmental delay and was a placid infant. He sat at 8 months, crawled at 14 months and walked at 2 years 11 months, after a long period of knee-walking. He vocalised complex babble at 8 months but this regressed and occurred intermittently. By 5 years, he was non-verbal and had no signs. He could follow a single command and was extremely hyperactive with impulsivity and a fascination with water. He has never had a seizure; an awake and sleep EEG at 10 months of age was normal. Trio exome sequencing identified a *de novo* heterozygous *KCNQ2* pathogenic variant (c.949G>A; p.A317T). *In silico* predictions (SIFT, Polyphen-2, MutationTaster) were consistent with a deleterious effect. This variant was absent from control populations in gnomAD (https://gnomad.broadinstitute.org/).

*Patient 2* was identified in a cohort study of individuals with DEEs with spike-wave activation in sleep (DEE-SWAS). ^33,35^ This patient (case 6) was a 4 years and 4 months old boy who presented with infantile epileptic spasms syndrome and hypsarrhythmia at 5 months of age (controlled by adrenocorticotrophic hormone from 10 months of age). At 1.8 years, Rolandic epileptiform discharges were seen which evolved to SWAS at 2.3 years. There was global developmental delay from birth, with regression after seizure onset, resulting in language delay and impairment of fine motor skills. Clinical gene panel identified a *de novo KCNQ2* variant (c.952C>G; p.L318V). *In silico* predictions (SIFT, Polyphen-2, MutationTaster) were consistent with a deleterious effect, and the variant was classified as pathogenic according to ACMG/AMP criteria).^36^

*Patient 3* is a female identified in a cohort of 4,293 individuals with severe undiagnosed developmental disorders, recruited as part of the Deciphering Developmental Disorders study via the genetics services of the UK National Health Service and the Republic of Ireland (patient ID DDD4K.02558). She carried the c.1066C>T; p.A356T variant in *KCNQ3*. Human Phenotype Ontology terms reported for this patient are: abnormal repetitive mannerisms; bimanual synkinesia; delayed speech and language development; microcephaly; moderate global developmental delay; poor motor coordination. No epilepsy at any age is reported, and no additional clinical detail is available for this patient. ^34^

### Homomeric or heteromeric Kv7.2 channels carrying pathogenic variants in the pore AG display a strong GoF *in vitro* phenotype

The A317T and the L318V variants found in patients 1 and 2, respectively, affect two consecutive residues located at the distal end of the S_6_ segment of Kv7.2 subunits, falling within the pore AG, a highly-conserved region undergoing large structural rearrangements during pore opening (**Fig. 1a,b**).^14,15,37,38^ To evaluate the functional consequences of these two variants, CHO cells were transfected with cDNAs plasmids encoding for Kv7.2, Kv7.2 A317T or Kv7.2 L318V subunits and the resulting currents were recorded with the whole-cell configuration of the patch-clamp technique. As shown in **Fig. 1c**, when depolarized from −140/-120 mV to 20 mV in 10 mV incremental steps, CHO cells expressing Kv7.2 channels generated voltage-dependent K^+^-selective currents with an activation threshold around −50/-40 mV and rather slow time-course of activation and deactivation. The half-activation potential (V_½_) and the slope of the I/V relationship (k) calculated from equation 1 (see Methods) were −20.13±1.09 mV and 12.03 ± 0.49 mV/e-fold, respectively (**Table 1**). At the holding voltage of −80 mV, Kv7.2 channels were closed and the ratio between the currents measured at the beginning of the depolarization step and those at the end of the 0-mV depolarization was close to 0 (**Table 1**).

**Fig. 1:**
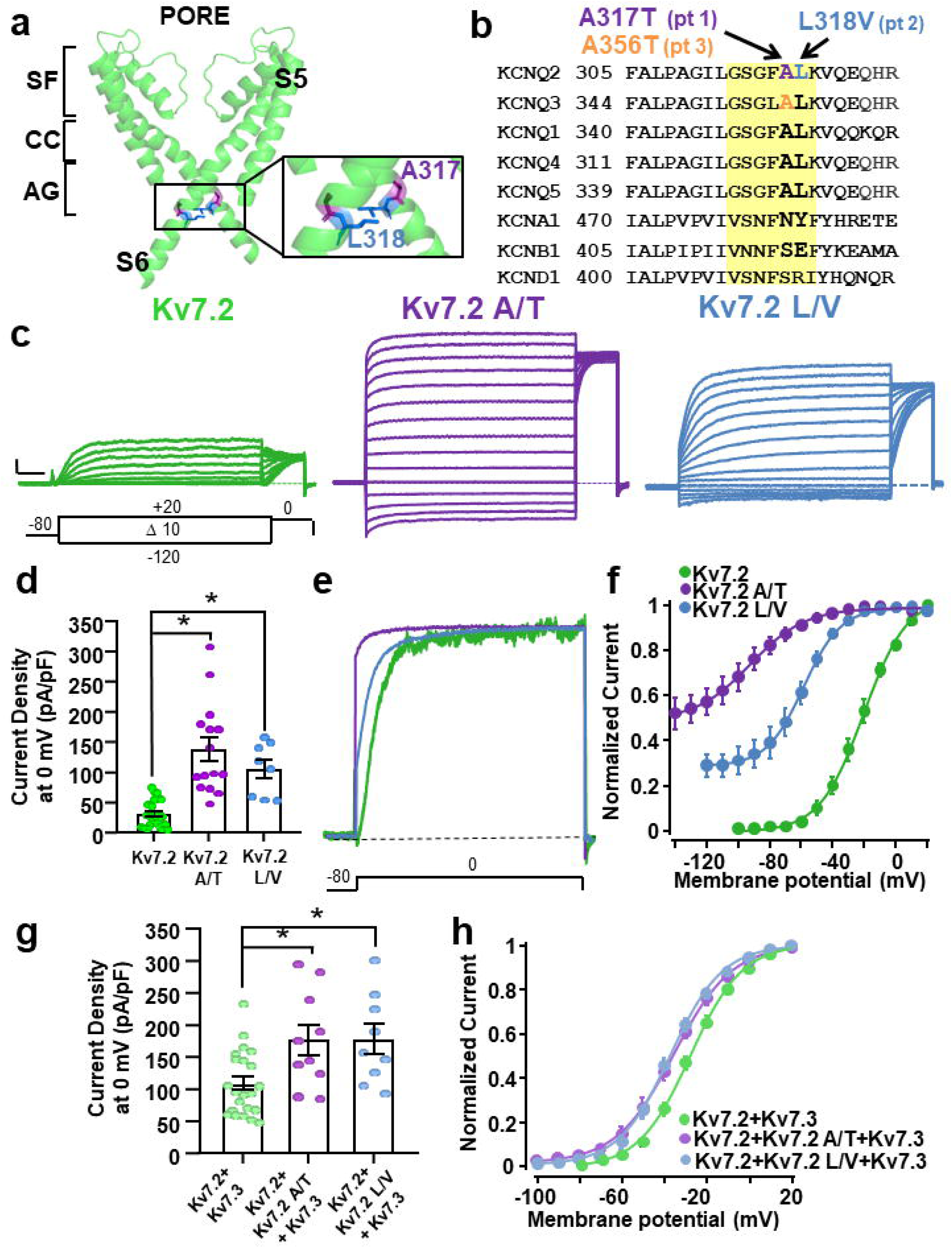
Biophysical properties of the K^+^ currents from Kv7.2 channels carrying variants in the pore AG. **a.** Kv7.2 structure; only the PD from two opposite subunits is shown for clarity. The enlargement highlights the A317 (purple) and the L318 (blue) residues. Selectivity filter (SF), central cavity (CC) and activation gate (AG) regions are indicated. **b.** Sequence alignment for the indicated Kv subunits in the S_6_ region; the investigated variants are highlighted. The region in yellow indicates the pore activation gate (AG). **c.** Macroscopic whole-cell currents from Kv7.2 (green), Kv7.2 A317T (purple), and Kv7.2 L318V (blue) homomeric channels in response to the indicated voltage protocol. The current scale is 200 pA, and the time scale is 200 ms. **d.** Quantification of the current densities recorded from cells transfected with the indicated cDNA constructs. * = p<0.05. **e.** Superimposed normalized current traces from Kv7.2, Kv7.2 A317T, and Kv7.2 L318V homomeric channels. Note the instantaneous current component in Kv7.2 A317T (purple), and Kv7.2 L318V channels at the beginning of the pulse. **f.** Conductance/voltage curves for the indicated channels; continuous lines represent Boltzmann fits of the experimental data to equation 1 in Methods. **g.** Quantification of current densities recorded from cells transfected with the indicated cDNA constructs. * = p<0.05. **h.** Conductance/voltage curves for the indicated channels; continuous lines represent Boltzmann fits of the experimental data to equation 1 in Methods.

**Table 1.** Biophysical properties of Kv7.2 and Kv7.3 mutant channels.

|  | **cDNA**<br>**trasfection (μg)** | **n** | **pA/pF at 0 mV** | **V<sub>1/2</sub> (mV)** | **k** | **l<sub>inst</sub>/l<sub>ss</sub>** | **TEA block (%)** |
| --- | --- | --- | --- | --- | --- | --- | --- |
| **Non-transfected** |  | 5 | 0.50 ± 0.20 | - | - |  |  |
| **Kv7.2** | 3 | 25 | 31.44 ± 4.57 | -20.13 ± 1.09 | 12.03 ± 0.49 | 0.03 ± 0.01 | 55.4 ± 7.6 (0.3 mM. n=3) |
| **Kv7.2 A317T** | 3 | 15 | 138.73 ± 20.30* | -92.93 ± 3.60* | 13.60 ± 1.92 | 0.81 ± 0.04* | 56.2 ± 3.4 (0.3 mM n=7) |
| **Kv7.2 L318V** | 3 | 8 | 105.26 ± 16.64* | -58.29 ± 1.28* | 11.94 ± 11.12 | 0.33 ± 0.03* |  |
| **Kv7.2+Kv7.3** | 1.5+1.5 | 20 | 109.08 ± 11.94 | -27.38 ± 1.23 | 11.38 ± 0.61 | 0.01 ± 0.003 | 50.8 ± 3.2 (3 mM n=9) |
| **Kv7.2+Kv7.2 A317T+Kv7.3** | 0.75+0.75+1.5 | 12 | 177.99 ± 25.04† | -36.74 ± 2.09† | 13.09 ± 2.09 | 0.12 ± 0.03† | 54.7 ± 2.2 (3 mM n=3) |
| **Kv7.2+Kv7.2 L318V+Kv7.3** | 0.75+0.75+1.5 | 9 | 178.09 ± 26.27† | -37.51 ± 1.40† | 11.98 ± 0.78 | 0.04 ± 0.01 | 50.25 ± 5.83 (3 mM n=3) |
| **Kv7.3** | 3 | 8 | 25.08 ± 5.94 | -31.08 ± 1.98 | 9.46 ± 1.17 |  |  |
| **Kv7.3 A356T** | 3 | 8 | 0.42 ± 0.12‡ | - | - |  |  |
| **Kv7.3+pcDNA3.1** | 1.5+1.5 | 6 | 24.68 ± 7.84 | -34.37 ± 2.34 | 6.75 ± 0.66 |  |  |
| **Kv7.3+Kv7.3 A356T** | 1.5+1.5 | 8 | 7.88 ± 2.68‡ | -34.61 ± 1.23 | 6.54 ± 0.67 |  |  |
| **Kv7.2+Kv7.3** | 1.5+1.5 | 20 | 109.08 ± 11.94 | -27.38 ± 1.23 | 11.38 ± 0.61 | 0.01 ± 0.003 | 50.77 ± 3.19 (3 mM n=9) |
| **Kv7.2+Kv7.3 A356T** | 1.5+1.5 | 9 | 371.91 ± 64.95† | -41.64 ± 3.26† | 11.98 ± 0.66 | 0.08 ± 0.03† | 51.87 ± 4.10 (3 mM n=5) |
| **Kv7.2+Kv7.3+Kv7.3 A356T** | 1.5+0.75+0.75 | 9 | 261.60 ± 50.25† | -44.02 ± 1.21† | 11.77 ± 1.23 | 0.05 ± 0.01† | 49.25 ± 3.06 (3 mM n=7) |

\* $P < 0.05$ versus Kv7.2.
† $P < 0.05$ versus Kv7.2 + Kv7.3.
‡ $P < 0.05$ versus Kv7.3.

When compared to Kv7.2 channels, both homomeric Kv7.2 A317T and L318V channels showed a significant 3-4-fold increase in the maximal current density (**Fig. 1c,d**; **Table 1**). In both mutants, the activation process revealed two clearly distinct kinetic components: an instantaneous voltage-independent component, followed by a slower voltage- and time-dependent component. At 0 mV, the instantaneous voltage-independent fraction accounted for 81±4% and 33±3% of the total current in Kv7.2 A317T and Kv7.2 L318V channels, respectively (**Fig. 1e**; **Table 1**). The V_½_ of the voltage-dependent component in Kv7.2 A317T and Kv7.2 L318V channels was hyperpolarized by 70 mV and 40 mV, respectively (**Fig. 1f**). Despite these dramatic changes in gating, both mutant channels retained their selectivity for K^+^ ions; indeed, the reversal potential of the currents from Kv7.2, Kv7.2 A317T and Kv7.2 L318V channels was close to that of a K^+^-selective pore (−74.2±0.65 mV for wild-type, WT, −75.8±0.96 mV for A317T and −73.8±1.81 mV for L318V; n=5 for each group; p>0.05 when compared to WT). Moreover, currents carried by mutant channels retained their sensitivity to the open-pore blocker tetraethylammonium (TEA)^39^ (**Table 1**), indicating that they were flowing through the central pore. Experiments in bi-ionic conditions with extracellular 15 mM K^+^, Rb^+^ or Cs^+^ ions revealed no difference in relative permeability between Kv7.2 and Kv7.2 A317T mutant channels (**Supplementary Fig. 1**).

In adult neurons, the M-current is mainly formed by Kv7.2/Kv7.3 heteromeric channels.^2^ To replicate *in vitro* the genetic status of patients 1 and 2 who were heterozygous for the *KCNQ2* pathogenic allele, functional studies were also carried out upon transfection of *KCNQ2* + *KCNQ2* A317T + *KCNQ3* and *KCNQ2* + *KCNQ2* L318V + *KCNQ3* cDNAs at a 0.5:0.5:1 cDNA ratio. Under these conditions, functional effects were similar to, but quantitatively smaller than, those observed in homomeric Kv7.2 channels, with a 50% increase in maximal current density (**Fig. 1g**) and a ∼10 mV hyperpolarizing shift in activation gating for both groups incorporating mutant subunits when compared to WT Kv7.2+Kv7.3 heteromeric channels (**Fig. 1h**). When compared to WT channels, no change in TEA sensitivity was observed in heteromeric channels incorporating mutant subunits (**Table 1**). Overall, these results suggest that channels carrying the A317T or the L318V variant display enhanced maximal current density, a hyperpolarized activation gating, and a reduced stability of the closed state; all these effects are consistent with an *in vitro* GoF phenotype. Notably, the A356 residue in Kv7.3 is paralogous to A317 in Kv7.2 (**Fig. 1b**), and the *KCNQ3* A356T variant, corresponding to A317T in *KCNQ2*, has been found in a DEE patient.^34^ In contrast to the strong *in vitro* GoF phenotype of homomeric Kv7.2 A317T channels, Kv7.3 A356T subunits did not form functional channels and reduced WT Kv7.3 currents when co-expressed (1:1 cDNA ratio) (**Supplementary Fig. 2a**; **Table 1**). However, a marked increase in current density and a ∼15 mV hyperpolarizing shift in activation gating was observed in heteromeric currents recorded from cells that were co-transfected with *KCNQ2* + *KCNQ3* A356T (1:1 cDNA ratio) or *KCNQ2* + *KCNQ3* + *KCNQ3* A356T (1:0.5:0.5 cDNA ratio) when compared to *KCNQ2* + *KCNQ3*-tranfected cells (1:1 cDNA ratio) (**Supplementary Fig. 2b,c**; **Table 1**). Thus, while showing LoF effects in homomeric or heteromeric configuration with Kv7.3 subunits, Kv7.3 A356T subunits, similarly to Kv7.2 subunits carrying the paralogous A317T variant, prompted strong GoF when expressed together with Kv7.2 subunits.

### Kv7.2 pore AG variants increase opening probability without affecting single channel currents or plasma membrane subunit expression

Both Kv7.2 A317T and Kv7.2 L318V variants caused a large increase in macroscopic current density, which depends on the number of functional channels (N), the single-channel current (i), and the channel opening probability (Po). To investigate variant-induced changes in each of these parameters, nonstationary noise analysis was performed. Mean currents and their respective variance were measured from Kv7.2-, Kv7.2 A317T-, and Kv7.2 L318V-expressing cells. When the variance was plotted against the average current and the data fitted with the parabolic equation 2 described in the Methods (**Fig. 2a**), no differences in the number of functional channels or in the single channel current (**Fig. 2b,c**) between Kv7.2 and Kv7.2 L318V channels were observed. By contrast, when compared to Kv7.2, Kv7.2 L318V channels showed a 2.5/3-fold increase in maximal Po (**Fig. 2d**). These data suggest that the Kv7.2 L318V mutant channels caused a strong GoF phenotype mainly through the increase of channel opening probability, without changes in the number of functional channels or their unitary conductance. While the data from Kv7.2 and Kv7.2 L318V channels allowed a detailed exploration of the entire open probability space, a similar approach in Kv7.2 A317T channels was limited by the small time-dependent component observed for the currents carried by these channel, leading to a less rigorous definition of the single channel estimates^40^ (**Supplementary Fig. 3a).** Nevertheless, unrestrained fitting of the available data revealed that, when compared to Kv7.2, Kv7.2 A317T channels, similarly to L318V channels, also displayed an increased opening probability with no changes in conductance or number (**Supplementary Fig. 3b-d**).

**Fig. 2:**
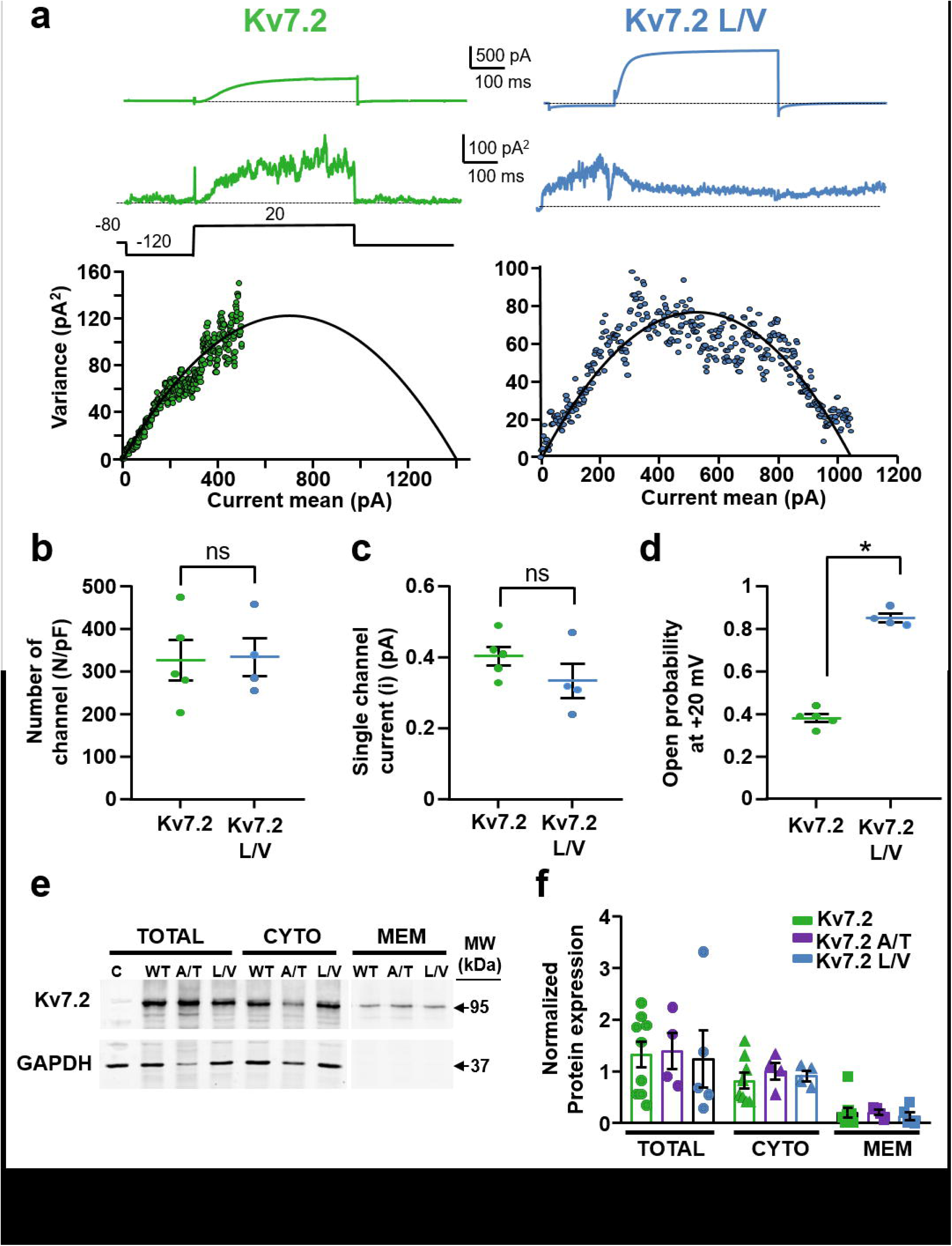
Non-stationary noise analysis of the K^+^ currents from Kv7.2 channels carrying variants in the pore AG. **a.** Representative average response to 100 pulses at +20 mV (top traces), variance (middle traces), and variance versus current mean plot (bottom traces) from Kv7.2 and Kv7.2 L318V, as indicated. The continuous lines in the variance/mean plots are parabolic fits of the experimental data to equation 2 in Methods. **b-d.** Quantification of the number of channels divided by capacitance (**b**), of the single-channel current (**c**), and of the opening probability at 20 mV (**d**) for the indicated channels. * = p<0.05. **e.** Representative Western-blot image from total, cytosol or plasma membrane protein fractions from CHO cells transfected with pcDNA3.1 (empty vector, C), Kv7.2 (WT), Kv7.2 A317T (A/T), or Kv7.2 L318V (L/V) subunits. On the right, the position of the estimated molecular mass for Kv7.2 (95 kDa) and GAPDH (37kDa) bands is shown. **f.** Densitometric quantification of 95 kDa band intensity for the indicated experimental groups. Data are expressed as means ± SEM.

Consistent with noise analysis results, Western Blotting experiments revealed no quantitative difference in the expression of Kv7.2, Kv7.2 A317T and Kv7.2 L318V subunits in total, plasma membrane or cytosolic fractions isolated from transfected CHO cells (**Fig. 2e,f**). In addition, when compared to WT Kv7.3, a slight increase in plasma membrane expression of Kv7.3 A356T mutant subunits was observed (**Supplementary Fig. 2d**), suggesting that the lack of functional currents of homomeric Kv7.3 A356T channels is not attributable to a mutation-induced impairment in membrane targeting and expression of mutant subunits.

### Currents from Kv7.2 pore AG variants are resistant to PIP_2_-dependent modulation

Channel opening of all Kv7 channels relies on the plasma membrane abundance PIP_2_; PIP_2_ is required for coupling VSD movement to PD opening of Kv7 channels during electromechanical coupling, and depleting PIP_2_ results in a closed pore with normal VSD activation.^41^ Several PIP_2_binding regions have been described in Kv7 channels, including the bottom part of the S_6_ segment.^13,38,42^ In WT Kv7.2 channels, increased PIP_2_ availability causes GoF consequences similar to those described above for the DEE variants affecting the AG, including a strong increase in maximal conductance and a hyperpolarizing shift in activation gating.^43,44^ To investigate the sensitivity of the Kv7.2 A317T and Kv7.2 L318V currents to PIP_2_ modulation, PIP_2_ levels were either increased by adding the water soluble PIP_2_ analogue dic8-PIP_2_ to the intracellular solution,^43^ or decreased by co-expressing a voltage-sensitive phosphatase from *Danio rerio* (DrVSP) which transiently reduces intracellular PIP_2_ levels when activated by strong depolarizations.^45^

Increasing cellular PIP_2_ levels by Dic8-PIP_2_ led to a progressive, time-dependent enhancement in Kv7.2 currents, together with a negative shift in their half-activation potential. By contrast, pipette inclusion of Dic8-PIP_2_ failed to affect current density in Kv7.2 A317T- and Kv7.2 L318V-transfected cells, and no shift in current half-activation potential occurred in those two mutants (**Fig. 3a-c**). Moreover, in CHO cells expressing Dr-VSP, depolarization to 100 mV time-dependently reduced Kv7.2 currents, reaching a maximum of 80% after 1 s; by contrast, currents carried by Kv7.2 A317T and Kv7.2 L318V channels were completely insensitive to DrVSP-mediated inhibition at all investigated time points (**Fig. 3d,e**). These data suggest that Kv7.2 channels carrying either A317T or L318V variant are fully insensitive to PIP_2_-dependent regulation.

**Figure 3:**
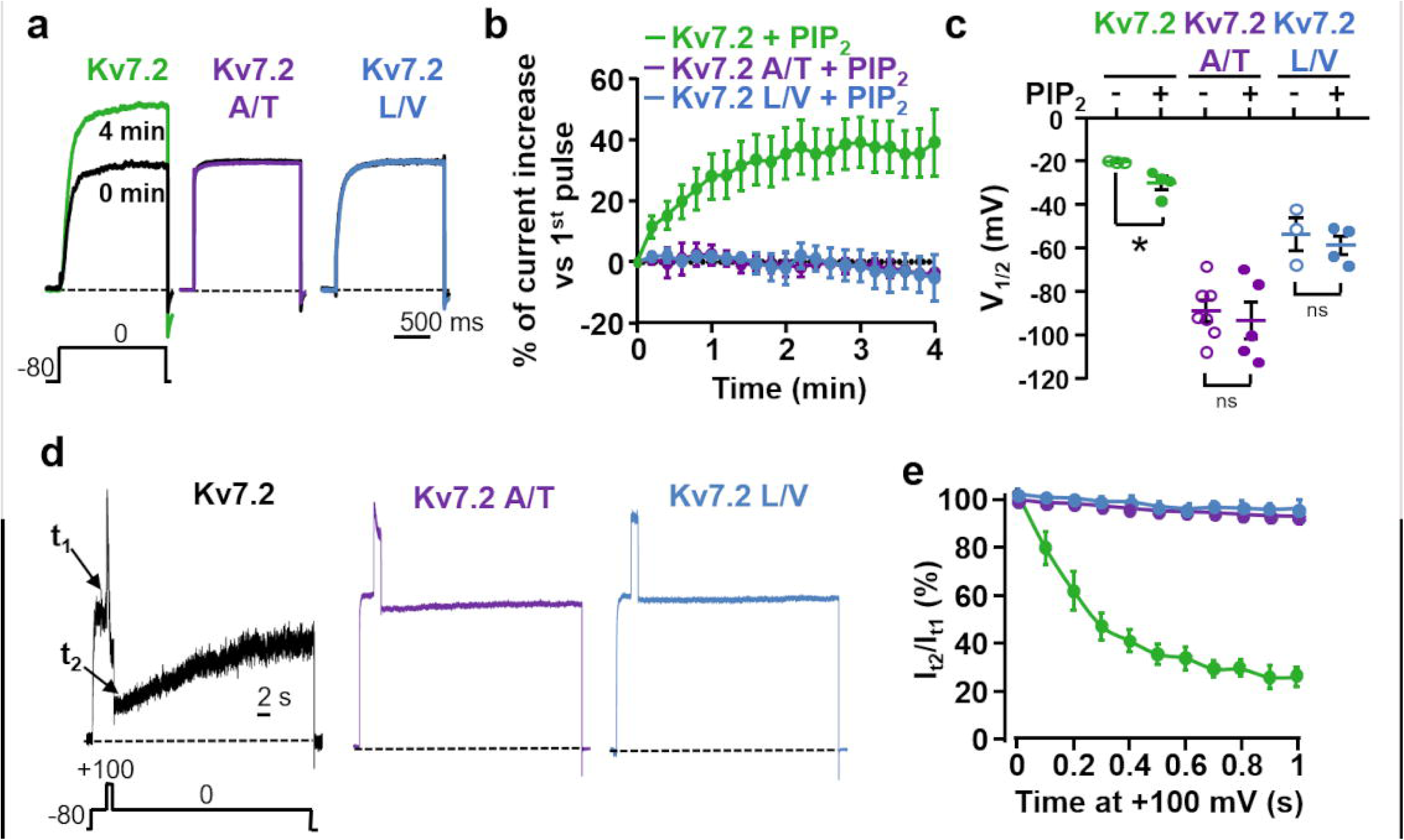
Effect of PIP_2_ level manipulations on Kv7.2, Kv7.2 A317T and Kv7.2 L318V currents. **a.** Macroscopic currents recorded in presence of 100 µM Dic8-PIP_2_ in the intracellular pipette solution at 0 min (immediately after patch rupture), and after 4 min of whole-cell intracellular dialysis. **b.** Normalized whole-cell currents for the 3 channels indicated as a function of time; 100 µM Dic8-PIP_2_ is present in the pipette solution. **c.** V_½_ values comparison between cells recorded in the absence (-) or in the presence (+) of 100 µM Dic8-PIP_2_ in the intracellular solution, after 4 min of whole-cell intracellular dialysis. * = p<0.05. **d.** Currents recorded in response to the indicated voltage protocol in cells expressing DrVSP and Kv7.2, Kv7.2 A317T or Kv7.2 L318V channels. Time scale, 2 s. **e.** Time-dependent current changes in cells co-expressing the indicated channels and DrVSP. Data are expressed as the ratio between the current values recorded at 0 mV immediately after (t2) and before (t1) the Dr-VSP–activating +100 mV depolarizing step as a function of time.

### Molecular modelling and MD simulations of the Kv7.2 AG variants

To gain insight into the molecular mechanism underlying the GoF effect of the described Kv7.2 pathogenic variants, all-atom MD simulations were performed. In addition to A317T and L318V, we also modelled the Kv7.2 G313S variant, paralogous to a pathogenic Kv7.5 variant also causing strong GoF effects.^46^ Given the relevance of allosteric coupling between the AG and the SF regions in K^+^ channels, and the occurrence of DEE-causing mutations at the filter residue D282,^34,47,48^ we modelled the WT sequence with either a charged or a protonated neutral aspartic acid at this site. WT and mutant protein pores were simulated in two distinct conformations, each with a different degree of AG opening, quantified by cross-distances (CDs) between pairs of Cα or side-chain atoms from diagonally opposed subunits. Five distances were used in total, one characterizing the SF via the Cα atoms of G279 (d1), and four characterizing the AG, using the Cα atoms of G313 (d2) and A317 (d3), the Oγ atoms of S314 (d4), and the Cδ atoms of L318 (d5). The latter two residues form the main AG constrictions in closed Kv7.2 structures.^49,50^

The first conformation simulated is derived from the cryo-electron microscopy (Cryo-EM) structure of the human Kv7.2 (hKv7.2, PDB ID: 7CR0),^49^ and showed a conductive filter (d1=8.03 Å) and a constricted gate, with d2 = 11.67 Å, d3 = 13.69 Å, d4 = 4.00 Å and d5= 7.14 Å **(Supplementary Fig. 4)**. Since no experimental structure of Kv7.2 with an open AG was available when we started this project, the open Kv7.2 conformation was homology modeled using the open human Kv7.1 (hKv7.1, PDB ID: 6V01) as template,^38^ resulting in a conducting filter (d1=7.97Å) and a wide AG (d2=13.55 Å, d3=19.62 Å, d4=12.51 Å and d5=17.30 Å). This modelled structure is nearly identical to that of four cryo-EM hKv7.2 structures with open AG conformations recently released (PDB ID: 8J01, 8J02, 8W4U, 8IJK), each obtained in the presence of both small molecules activators and PIP_2_.^15,50^ In fact, the comparison resulted in root-mean-square deviations (RMSDs) between 0.331Å and 0.889Å, calculated over all 115 Cα atoms of the simulated chains, and very similar values of CDs (**Supplementary Fig. 5**).

The channels were embedded in explicit POPC membrane bilayers, solvated with water and ions, and simulated in five independent replicates for both closed and open AG conformations, obtaining a cumulative time length of 24.5 μs. All protein structures were stable during the MD simulations and did not substantially deviate from the starting conformations, as evidenced by backbone RMSDs averaged over the replicas that are stationary around values between 2.5 and 3 Å (see **Supplementary Figs. 6** and **7** for closed systems and **Supplementary Figs. 8** and **9** for open systems).

To investigate the impact of the mutations on the protein structure, we first looked at the size of the channel pore along the simulated trajectories. As for the WT channel, the open and closed pore radius profiles, averaged over the replicas, superimposed to those of the corresponding experimental structures, confirming that the pore geometry is maintained over the simulations, even with neutral D282 (**Supplementary Fig. 10**). When we compared the pore profiles of A317T, L318V and G313S to those of the WT Kv7.2, the results for closed and open AG structures differed. In particular, the A317T mutation induced a widening of the closed AG, from above the level of G313 down to the cytoplasm, that was highly pronounced in correspondence of the mutated residue (**Figs. 4a,b**). Interestingly, we observed a significant loosening of the two distinctive closed-gate constrictions at residues S314 and L318. This localized broadening of the pore makes it more accessible to water, as shown by the average number of water molecules computed along the channel axis (**Fig. 4c**). The difference is most noticeable in the CC region, particularly around V302, where the WT system is essentially dry, while A317T accommodates about eight water molecules. Both features are consistent with a GoF, since a broadened gate could facilitate the transition of the mutated channel towards the open state, and the concomitant increased hydration of the cavity may promote ion permeation.^12^ In contrast, the WT and A317T open structures trajectories showed no difference in pore profiles (**Figs. 4d,e**) or extent of hydration of the CC (**Fig. 4f**). In line with the enhanced hydrophilicity of the substitution, more water molecules entered the cytoplasmic side of the mutated pore, at A317T and below, but the difference remained limited to that region.

**Figure 4:**
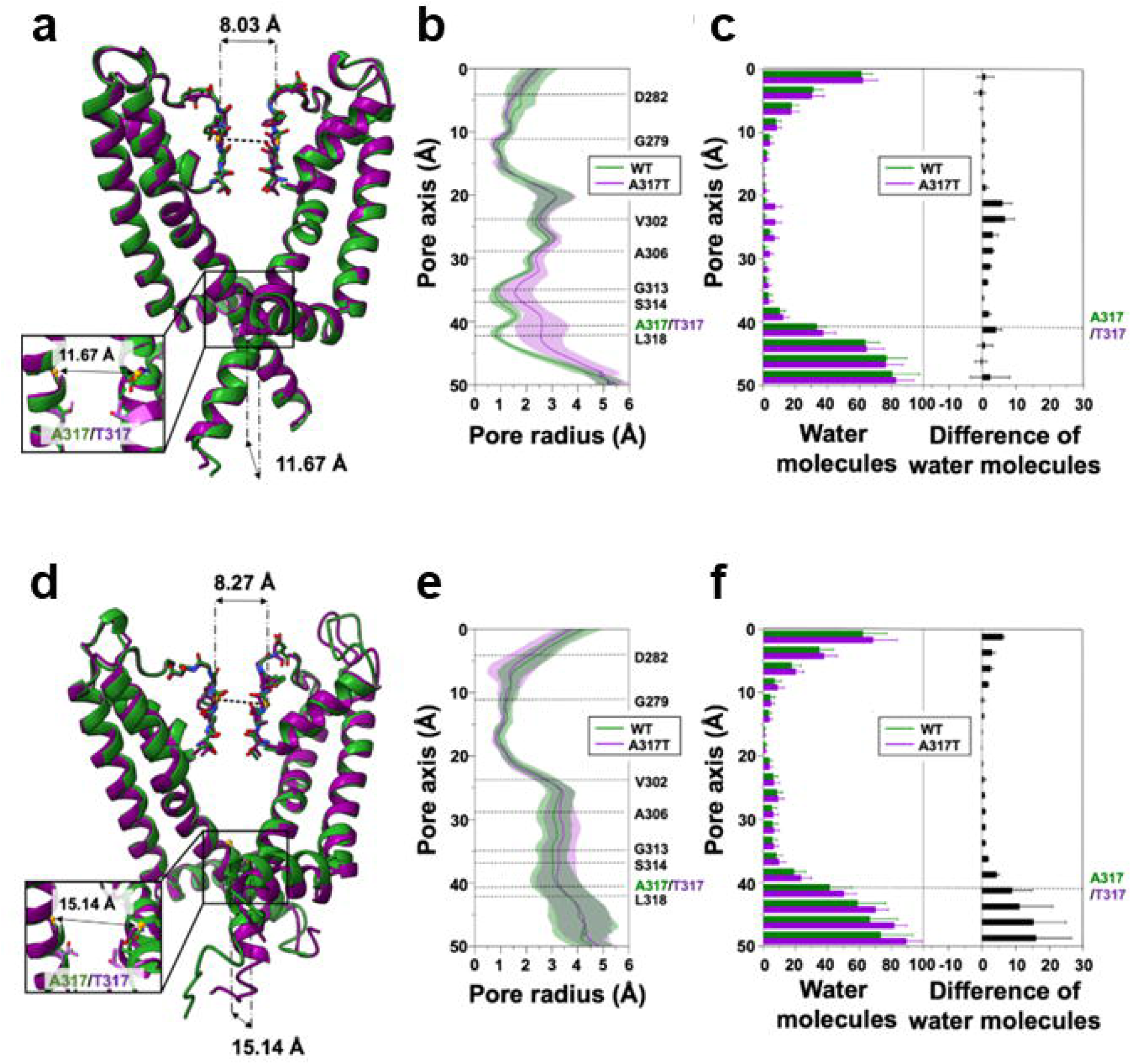
Molecular Dynamics simulations of the A317T Kv7.2 variant. **a.** Superposition of WT (green) and A317T (purple) representative Kv7.2 closed structures after equilibration and before MD production simulations. **b.** Channel radius profiles along the pore axis of the two proteins, averaged over all simulated replicas. Shaded regions indicate standard deviations. **c.** Distribution of water molecules along the channel axis. Average and standard deviations are calculated over all replicas. **d-f**. Same as **(a-c),** but for the open structures.

Given the rearrangement of the closed A317T AG into a wider conformation, we searched for interactions involving the mutated amino acid that could be responsible for its stabilization. We found that the inserted threonine sidechains can form hydrogen bonds (HBs) with S314 from adjacent subunits (average persistence times of 54.8%, 68.0%, 64.3%, and 45.1% for the four pairs of chains, **Supplementary Fig. 11**), which cause the serine sidechains to point away from the pore lumen, removing the stable constriction they form in the WT closed conformation (**Figs. 5a,b)**. Notably, the new S314 orientation is similar to that observed in a recently determined open Kv7.2 structure, bound to the Ebio1 activator^50^ (**Supplementary Fig. 12**). In summary, the substituted T317 is directly involved in inter-chain bonds that cause the partial widening of the closed AG, generating a conformation that allows more water entry and may be more prone to open completely. Conversely, in the open conformations, the chains are at a larger distance from each other, the intersubunit T317-S314 interaction between them is absent, and the mutated threonine mostly connects to G313 from the same subunit **(Figs. 5c,d; Supplementary Fig. 11)**.

**Figure 5:**
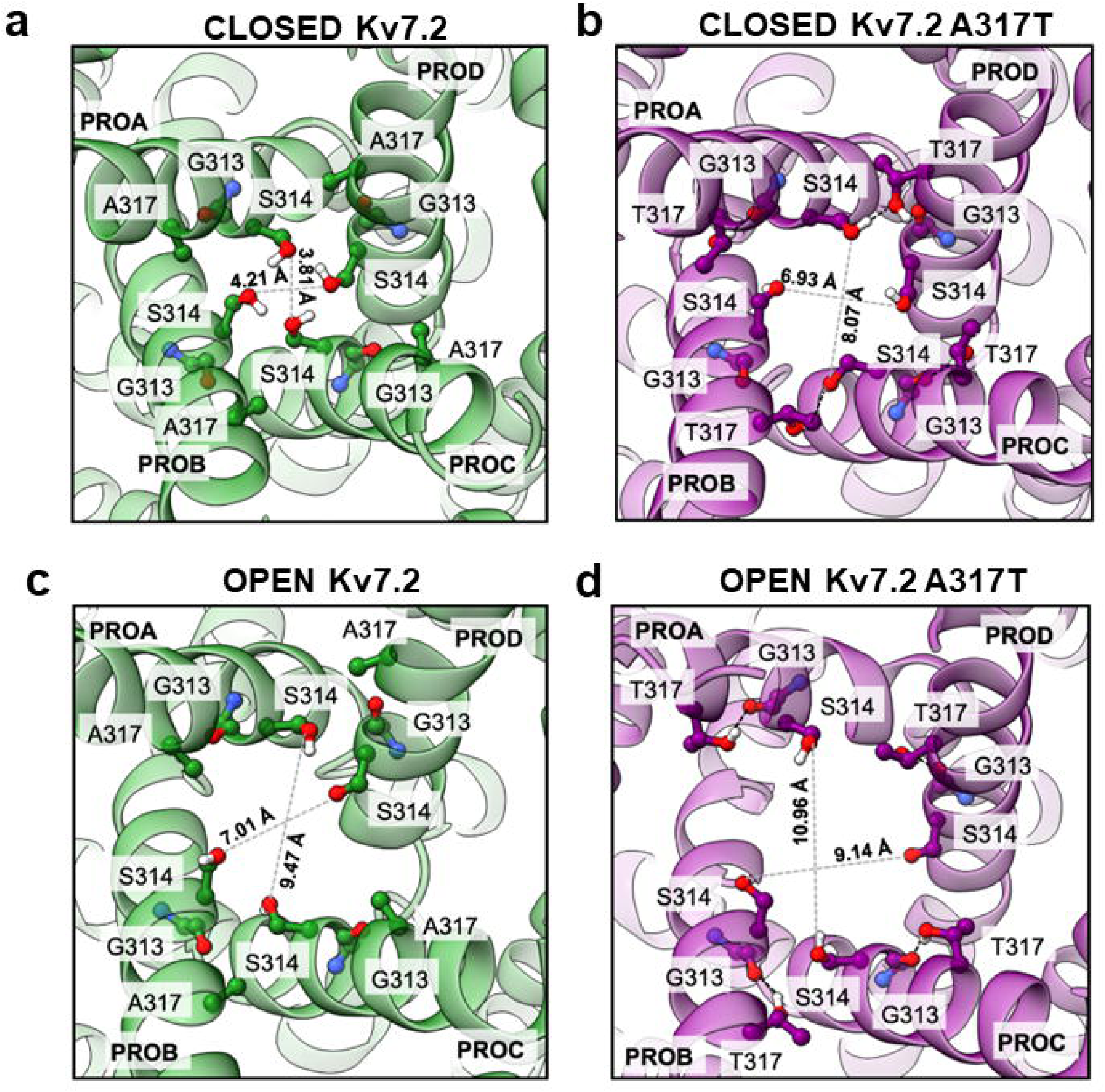
Closed-up cytosolic views of AG conformations from WT and A317T Kv7.2 channel simulations. **a.** Closed WT. **b.** Closed A317T variant. **c.** Open WT. **d.** Open A317T variant. Cross distances are indicated as gray dashed line. Where present, black dashed lines indicate HBs.

To verify that the structural effects observed in the A317T channel simulations do not depend on the membrane model, we performed additional calculations for the mutated closed pore embedded in bilayers of different lipid composition (pure POPE, POPG, DMPC, and a mixture POPC-CHOL-PIP2). In all cases, we observed a broadened AG and an enhanced hydration of the CC, comparable with those found with the pure POPC bilayer (**Supplementary Figs. 13** and **14)**.

As for the L318V variant, we also found a broadening of the AG region in the closed conformation, although to a smaller extent than in A317T (**Figs. 6a,b**). In L318V, the widening is mostly pronounced at the mutated residue, corresponding to the lower constriction site. Similarly to A317T, widening of the closed AG results in an enhanced hydration of the central cavity (**Fig. 6c**). Moreover, the WT and L318V open structures trajectories showed no difference in pore profiles (**Figs. 6d,e**) or hydration of the CC (**Fig. 6f**).

**Figure 6:**
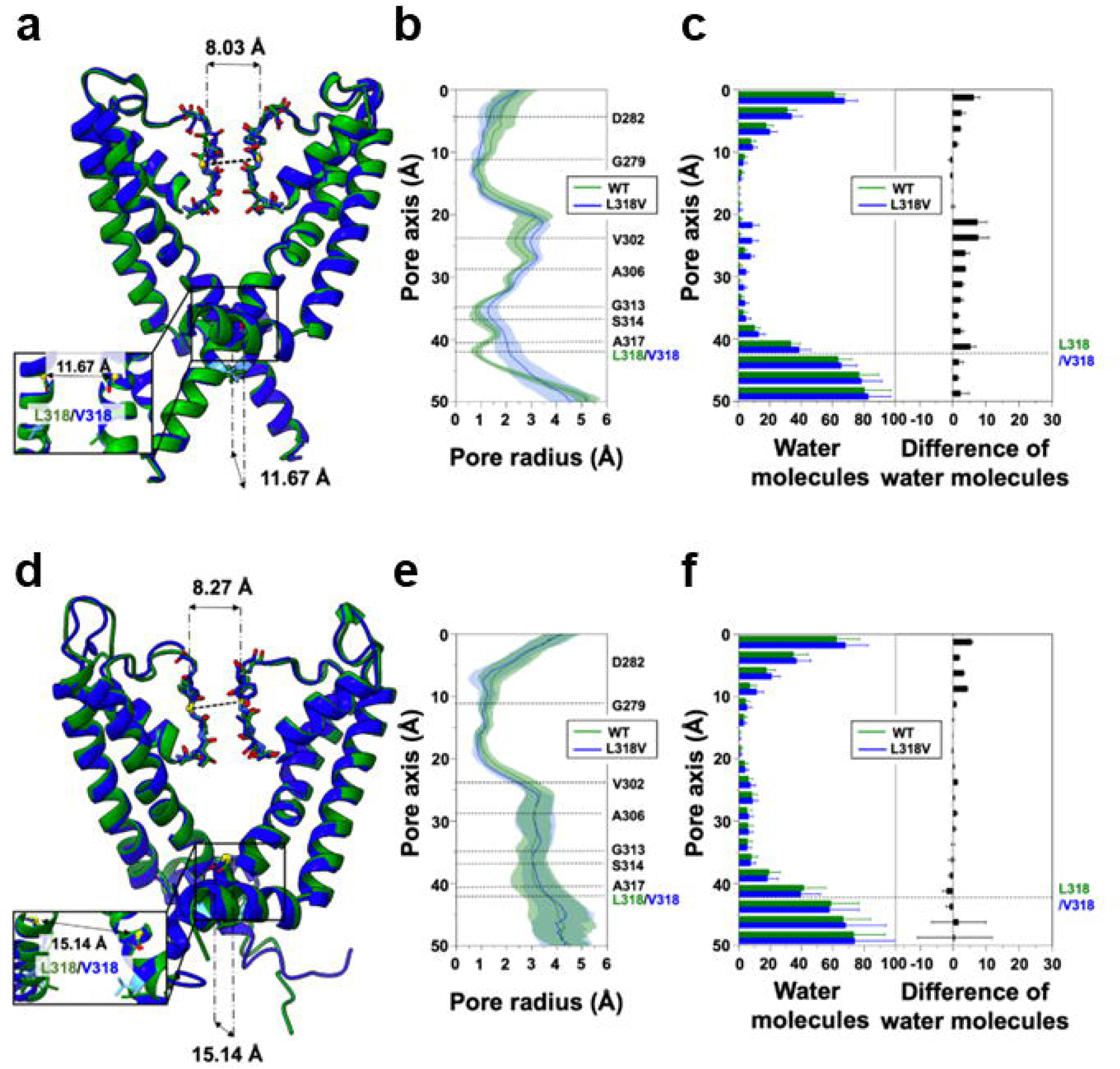
MD simulations of the L318V Kv7.2 variant. **a.** Superposition of WT (green) and L318V (blue) representative Kv7.2 closed structures after equilibration and before MD production simulations. **b.** Channel radius profiles along the pore axis of the two proteins averaged over all simulated replicas. Shaded regions indicate standard deviations. **c.** Distribution of water molecules along the channel axis. Average and standard deviations are calculated over all replicas. **d-f.** Same as **(a-c),** but for the open structures.

To further investigate the structural consequences associated with GoF Kv7.2 variants, we also simulated the non-pathogenic G313S variant, which corresponds to the DEE-causing G347S variant in Kv7.5; similarly to the A317T and the L318V variants investigated here, the Kv7.2 G313S also induces GoF effects by increasing the single channel open probability.^46^ The results obtained were remarkably similar those of A317T (**Supplementary Fig.15**): the mutation induced a widening of the closed AG (at the level of G313 and A317) and, in this case, also of the CC (around V302), associated with an increased hydration of this region. Conversely, in the open conformation, no major difference is detected in pore size or CC hydration. A comparison between four representative Kv7.2 snapshots of the WT, G313S, A317T, and L318V closed AG extracted from the respective simulations is reported in **Supplementary Fig. 16**, highlighting the enhanced pore opening and cavity hydration of the variants.

To further characterize the WT and mutant channel structures, we looked at the time-evolution of the cross-distances at the level of the SF (d1) and the AG (d2, d3, d4 and d5). All closed AG structures showed d1 values around 8 Å, generally considered indicative of a conductive conformation (**Supplementary Fig. 17)**. Smaller fluctuations and a preponderance of less constricted (i.e., conducting) filter structures are observed in the mutants (A317T and L318 in particular). As for the AG distances, closed A317T and G313S display higher and more fluctuating d2 and d3 values than the WT (**Supplementary Figs. 18** and **19**), while L318V shows a less clear signal. Conversely, d4 and d5 clearly indicate a widening of the closed AG constrictions in all three variants, consistent with the measurements of pore profiles (**Supplementary Figs. 20** and **21**, respectively). Finally, in the open AG simulations, no significant differences are present between WT and mutants in terms of cross-distances (**Supplementary Figs. 22-24**).

## DISCUSSION

*In vitro* functional features of disease-causing ion channel variants are often associated to specific clinical traits, such as disease severity, comorbidities, and therapy response. Thus, defining the molecular patho-mechanism(s) responsible for variant-induced changes in ion channel function may unveil prognostic clues and allow patient stratification for personalized therapeutic approaches. In the present study, we describe pathogenic variants affecting residues in the pore AG of Kv7.2 leading to strong PIP_2_-independent GoF features with a marked hyperpolarizing shift of voltage-dependent gating and a significant increase in current density. The complementary MD simulations analysis revealed that the substituted residues widen the pore AG by promoting a complex network of intra- and inter-subunit interactions, thus providing a plausible atomistic explanation for the observed results.

### Phenotypic Spectrum and Genotype-Phenotype Correlations in KCNQ2/3-related diseases

Pathogenic variants in *KCNQ2* have been identified in patients with a broad spectrum of predominantly neonatal-onset epileptic phenotypes, including SLFNE at the mild end, to *KCNQ2* early-infantile DEE at the severe end.^2,3,5^ Additional rarer phenotypes include: 1. neonatal encephalopathy with non-epileptic myoclonus, hypotonia, perinatal respiratory failure, and profound developmental delay;^23^ 2. non-neonatal-onset DEE with infantile/childhood-onset epilepsy, including West syndrome;^20–22^ 3. isolated ID without epilepsy, with various degrees of motor and speech delay.^8^ Over the years, a correlation has been established between functional effects *in vitro* of *KCNQ2* variants and the clinical phenotypic spectrum, with variants associated with severe DEEs causing a marked (>50%) decrease in channel function, often with dominant-negative mechanisms,^17,18^ whereas variants found in most self-limiting forms causing haploinsufficiency and milder LoF (<50%) *in vitro*.^2^ In addition to LoF, few *KCNQ2* GoF variants have been described, mostly in patient with rarer phenotypes of developmental disorders with moderate to severe ID in the absence of neonatal seizures.^3^ Indeed, the phenotypic characteristics of the three patients described herein are consistent with this stratification, with GoF effects associated with ASD, global developmental delay and without seizures in patient 1, and delayed development with infantile epileptic spasms syndrome followed by DEE-SWAS in patient 2;^33^ neither patients had neonatal-onset seizures.

While hundreds of pathogenic variants have been reported in *KCNQ2*, a much lower number of individuals carrying pathogenic variants has been described in the closely-related *KCNQ3* gene, with phenotypes ranging from SLFNE to more severe cases with neurodevelopmental disorders with or without seizures.^4^ A similar correlation to that observed for *KCNQ2* pathogenic variants has been suggested for the *in vitro* functional effects of *KCNQ3* pathogenic variants and phenotypic spectrum, with variants in most SLFNE cases causing milder LoF effects, and variants associated with more severe phenotypes causing more dramatic LoF effects.^51,52^ Notably, also for *KCNQ3*, *de novo* missense pathogenic variants causing strong GoF effects^28^ have been described in children with global developmental delay (likely similar to that justifying inclusion of patient 3 in the NDD cohort),^34^ ASD, and frequent SWAS.^24^

### Changes in VSD stability or VSD-PD coupling underlie GoF effects by known variants causing DEEs in KCNQ2 and KCNQ3

Recent structural evidence in Kv7.1, ^38,53–56^, Kv7.2,^15,37^ and Kv7.4^14,57^ have revealed important clues regarding their voltage-dependent gating mechanism, as well as their regulation by CaM and PIP_2_, providing a framework for the interpretation of the molecular mechanism by which disease-causing variants affect channel function. Although some details are still unknown, and specific structural features differentiate Kv7s from other voltage-gated K^+^ channels,^11^ the overall mechanism for voltage-dependent gating can be summarized as follows. Membrane depolarization induces a vertical shift of the S_4_ segment of the VSD, with the first two positively-charged arginine residues (R1, R2) passing through the gating charge transfer center formed by a phenylalanine residue in S_2_ and two negatively charged residues in S_2_ and S_3_. This upward dislocation of S_4_ induces a lateral rotation and an upward motion of the S_4_–S_5_ linker which bends the S_6_ C-terminal half of an adjacent subunit and leads to the opening of the pore. In the closed pore state of Kv7.2 (PDB: 7CR0), the diagonal atom-to-atom distance is ∼4 Å at the constriction-lining residue S314. In contrast, in the open state, the constriction-lining residue is G310, which has a diagonal atom-to-atom distance of 8 Å. Pore opening is also accompanied, and possibly facilitated by the fact that, upon depolarization, the HA and HB helices in the proximal C-terminus, where CaM binds in the closed channel configuration, undergo an almost 180^°^ rotation, causing the HA and S_6_ helices to join in a continuous helix and CaM to be released from its attachment site in the S_2_-S_3_ linker.

With the exception of the G239S in S_5_,^21^ all described *KCNQ2* GoF variants affect the VSD of Kv7.2. For example, the recurrent variants at R2 in the S_4_ of Kv7.2 increased the maximal current density, also causing a marked hyperpolarizing shift in the voltage-dependence of activation, and an acceleration of activation kinetics, suggestive of an increased sensitivity of the opening process to voltage.^23^ Given that R2 forms an intricate network of electrostatic interactions with neighboring negatively-charged residues in the resting configuration of the VSD, while no interaction involving R2 occurs when the VSD occupies the active configuration, these GoF effects occur as a consequence of a mutation-induced destabilization of the resting VSD configuration, leading to constitutive channel activation and pore opening.^28^ A similar molecular mechanism likely explains the GoF effects *in vitro* caused by variants affecting other VSD residues in both KCNQ2^20,22^ and KCNQ3.^24,28^ Although no direct structural confirmation for such a mechanism is yet currently available, voltage-clamp fluorometry studies in both Kv7.2^29^ and Kv7.3^30^ confirm that variants affecting charged residues in the proximal part of S_4_ including R1 and R2 alter channel function by directly impacting S_4_ activation, whereas those in the distal part of S_4_ do not interfere with VSD movement during activation gating, but rather with VSD-PD coupling.

### Constitituive pore opening by KCNQ2/3 pathogenic variants in the AG

We describe the functional properties of two variants in *KCNQ2* causing developmental encephalopathy and affecting consecutive residues in the S_6_ pore AG of Kv7.2 channels, namely A317T and L318V. When expressed as homomers or heteromers, both variants displayed a GoF *in vitro* phenotype, with a dramatic increase in current density and a marked facilitation of the voltage-dependent opening process. Non-stationary noise analysis revealed that these changes, highly consistent with a GoF in vitro phenotype, were due to an increase in the channel open probability, with no changes in the number of functional channels or their single-channel current amplitude. MD simulations indicate that the A317T variant, which affects a highly flexible region of S_6_ where pore widens as a consequence of the mechanical forces imposed by VSD repositioning during activation, stabilizes a broadened pore configuration in the closed channel. In particular, the formation of hydrogen bonds between the −OH group of T317 and the side chain of S314 from an adjacent subunit removes AG constrictions, inducing a constitutive enlargement of its diameter. Such stabilization of the pore in an open-like conformation, which occurs independently from voltage-dependent VSD repositioning or VSD-PD coupling, is observed also in simulations of L318V and G313S, and is consistent with the described GoF features of channels incorporating those Kv7.2 mutant subunits. Notably, Kv7.3 subunits carrying the A356T variant, corresponding to A317T in Kv7.2 and also causing neurodevelopmental delay,^34^ induced strong GoF effect, although only when expressed with Kv7.2 or Kv7.2 and Kv7.3 s subunits. A similar lack of functional currents in homomeric state, but a strong GoF when co-expressed with Kv7.2 subunits, has been previously reported for another KCNQ3 variant affecting the turret domain in the S_5_-S_6_ linker (Kv7.3 A315V).^58^

### Pore opening by Kv7.2 pore AG GoF variants is PIP_2_-independent

The endogenous lipid PIP_2_ is essential to the activation of KCNQ channels.^13,59^ Although PIP_2_-bound structures of Kv7.1^54^, Kv7.2,^15^ and Kv7.4^57^ channels have been recently described, the PIP_2_ activation mechanism is still unclear.^11^ PIP_2_ seems to facilitate voltage-dependent pore opening by binding to the S_2_-S_3_ linker where it disrupts the (likely inhibitory) interaction between the VSD and CaM (site I in Kv7.1 and Kv7.4), and/or stabilizing the channel in the open-state by direct interactions with positively-charged residues in the distal S_6_ or in the proximal C-terminus (site II in Kv7.2 and Kv7.4).

The functional changes triggered by the herein-described variants at the pore AG in Kv7.2 and Kv7.3 channels are similar to those promoted by PIP_2_ binding. Currents from both Kv7.2 A317T and L318V channels are fully insensitive to changes in PIP_2_ availability, as mutant channels are resistant to inhibition or potentiation when PIP_2_ levels are decreased or increased, respectively. Intriguingly, the L351K substitution in Kv7.1, affecting the residue paralogous to L318 in Kv7.2, also destabilized the closed pore state and rendered the channels insensitive to PIP_2_ depletion.^13^ The fact that pore opening in Kv7.2 A317T and L318V channels is fully PIP_2_-independent allows us to hypothesize that these variants promote structural changes leading to a pore configuration which largely recapitulates the open pore structure promoted by PIP_2_ in WT Kv7.2 channels. Such hypothesis is largely confirmed by the identical permeability ratios for K^+^/ Rb^+^ or K^+^/Cs^+^ between wild-type Kv7.2 and Kv7.2 A317T channels, as well as by our MD simulations showing no marked difference in the average cross-distances among the open pores of Kv7.2 and Kv7.2 A317T and L318V channels.

### Conclusions

Investigating variant-induced changes in channel function reveals disease pathogenetic mechanisms and allow to establish correlations among genotype, clinical phenotype, and *in vitro* features which might be critical for diagnostic evaluation, prognostic predictions, and targeted therapeutic management. Here, we provide evidence showing that pathogenic variants in the pore AG of Kv7.2 (and Kv7.3) channels cause strong GoF by a novel mechanism, namely a VSD-independent stabilization of the open pore configuration, thereby expanding genotype-phenotype correlations and providing new hypotheses for Kv7 channel gating and regulation deserving structural testing. Our results also highlight a mechanism which could be exploited for the pharmacological treatment of neurodevelopmental disorders caused by neuronal Kv7 channel dysfunction. In fact, in close analogy to the Kv7.1 activator ML277^60^ which promotes antiarrhythmic and cardioprotective actions^61^ by a direct, PIP_2_-independent stabilization of the open pore AG,^55^ selective Kv7.2 activators have been recently shown to facilitate channel opening via a twist-to-open motions involving S_6_ residues in the pore AG,^49^ thus prompting structural rearrangements similar to those triggered by the herein-described variants.

## METHODS

### Patient Recruitment and Genetic Analyses

Patient 1 was recruited at the Epilepsy Research Centre, Department of Medicine, University of Melbourne, Australia. Detailed phenotyping and trio exome sequencing were performed. The patient had a *de novo KCNQ2* missense variant, A317T. The study was approved by the Human Research Ethics Committee, Austin Health. Written informed consent for study participation was obtained from his parents. The L318V variant in Kv7.2 (patient 2)^33^ and the A356T variant in Kv7.3 (patient 3)^34^ have been reported previously in large cohort studies of neurodevelopmental disorders and DEEs.

### Mutagenesis and Heterologous Expression of KCNQ2 and KCNQ3 cDNAs

Mutations were engineered in human *KCNQ2* and *KCNQ3* cloned into pcDNA3 by site-directed mutagenesis. Channel subunits were expressed in Chinese Hamster Ovary (CHO) cells by transient transfection. CHO cells were transfected using Lipofectamine 2000.^17^

### Whole-Cell Electrophysiology and Nonstationary Noise Analysis

Currents from CHO cells were recorded at room temperature (20-22 °C) 1-3 days after transfection. The extracellular solution contained (in mM): 138 NaCl, 5.4 KCl, 2 CaCl_2_, 1 MgCl_2_, 10 glucose, and 10 HEPES, pH 7.4 with NaOH. The intracellular solution contained (in mM): 140 KCl, 2 MgCl_2_, 10 EGTA, 10 HEPES, 5 Mg-ATP, pH 7.3-7.4 with KOH. To evaluate PIP_2_ sensitivity, the water soluble, synthetic PIP_2_ analogue dic8-PIP_2_ (100 μM) was added to the intracellular solution. The pCLAMP software (version 10.0.2) was used for data acquisition and analysis. Linear cell capacitance (C) and series-resistance (R_S_) calculations, as well as cell capacitance compensation, were performed as described.^62^ All currents were corrected offline for linear capacitance and leakage currents using standard subtraction routines (Clampfit module of pClamp 10). Current densities (expressed in pA/pF) were calculated as peak K^+^ currents divided by C. Data were acquired at 0.5-2 kHz and filtered at 1-5 kHz with the 4-pole lowpass Bessel filter of the amplifier. No corrections were made for liquid junction potentials. To generate conductance-voltage (G/V) curves, cells were held at −80 mV, then depolarized for 1.5 s from −80 mV to 20/80 mV in 10 mV increments, followed by an isopotential pulse at 0 mV of 300-ms duration; current values recorded at the beginning of the 0 mV pulse were measured, normalized and expressed as a function of the preceding voltages. The data were then fit to a Boltzmann distribution of the following equation 1:

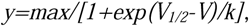

where V is the test potential, V_½_ the half-activation potential, and k the slope factor.^17^

Non-stationary noise analysis data were acquired and analysed as previously described.^46^ In short, 100 pulses of 500-ms duration were applied to a test voltage of 20 mV from the holding potential of −80 mV at 1 Hz frequency. To maximize the voltage-dependent component of the currents, a 100-ms prepulse at −120 mV was delivered before each 500-ms depolarisation to +20 mV. Currents were lowpass filtered at 10 kHz and sampled at 100 kHz. Variance was obtained by averaging the squared difference of consecutive records after appropriate scaling. Potential artifacts due to current rundown (<20%) were minimized by measuring the differences in the current on successive sweeps between consecutive pulses. Baseline variance was subtracted, and the variance-mean plot was fitted by the equation 2:

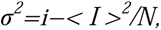

where σ^2^is the variance, < I > is the mean current, i is the single-channel current, and N is the number of channels. During fitting, i and N were not constrained, and points were not weighted by their standard errors (SE). The maximal open probability was calculated as P_max_= _max_/(Ni). TheGePulse and Ana programs (http://users.ge.ibf.cnr.it/pusch/programs-mik.htm) were used, respectively, for data acquisition and analysis.

Permeability to monovalent cations was measured as previously described.^63^ Briefly, an extracellular solution contained (in mM) 127.4 NaCl, 15 KCl, 2 CaCl_2_, 1 MgCl_2_, 10 glucose, and 10 HEPES, pH 7.4 with NaOH was used to measure K^+^ permeability (P_K_). KCl was replaced by 15 mM RbCl or CsCl to estimate Rb^+^ and Cs^+^ permeabilities. Currents were activated by 500-ms pulses at 20 mV and deactivated by 1500 ms steps from −120 mV to −30 mV to estimate their reversal potential under each bi-ionic condition. The outer monovalent cations’ permeability relative to K^+^ (P_X_/P_K_) was determined using the Goldman-Hodgkin-Katz equation: P_X_/P_K_ = ([K]_o_/[X]_o_)exp – (FΔE_rev_/RT), where ΔE_rev_ is the shift in reversal potential when K^+^ is replaced by the test cation X. The constants F, R, and T in the equation also adhere to their conventional thermodynamic meanings.

### Cell-Surface Biotinylation and Western Blot

Plasma membrane expression of wild-type and mutant Kv7.2 or Kv7.3 subunits in CHO cells was investigated by surface biotinylation of membrane proteins in transfected cells one day after transfection as described.^28^ Briefly, cells were incubated with Sulfo-NHS-LC-Biotin (0.5 mg/mL; Thermo Fisher, Monza, Italy), a cell-membrane impermeable reagent, for 20 min at 4 °C; the reaction was quenched with 0.1 M glycine in phosphate-buffered saline (PBS) pH 8.0 (3 washes, 20 min each). Subsequently, cells were lysed and protein lysates were incubated with streptavidin beads (Thermo Fisher, Monza, Italy) to isolate biotinylated proteins. Channel subunits in streptavidin precipitates and total lysates were analysed by Western blotting on 8% sodium dodecylsulfate polyacrylamide gel electrophoresis (SDS-PAGE) gels and transferred onto a polyvinylidene fluoride membrane (Merck, Milan, Italy).

Membranes were incubated overnight at 4 °C with rabbit polyclonal anti-Kv7.2 (dilution 1:1000; GeneTex, Irvine, USA) or anti-Kv7.3 (dilution 1:1000; Alomone Lab., Jerusalem, Israel); secondary antibodies conjugated to the fluorescent-dyes StarBright Blue 520 or StarBright Blue 700 were incubated for 1 h at RT. An anti-GAPDH antibody (dilution 1:2000: ThermoFisher, Monza, Italy) was used to check for equal protein loading and to assess the purity of the plasma-membrane preparation. Blot images, as well as fluorescence signals were acquired with ChemiDoc™ Touch Imaging System (Biorad, Milan, Italy). Images were analysed by using Image Lab Software (version 6.1; Biorad, Milan, Italy). Intensities of Kv7.2 and Kv7.3 bands were normalized to that of GAPDH. Levels of plasma-membrane Kv7 protein were normalized to total Kv7 protein input.

### Statistics

Data are expressed either as means ± SEM for number of independent samples (n). Normal distribution was assessed using the D’Agostino–Pearson normality test. To compare two normally distributed sample groups, the unpaired two-tailed Student’s t-test was used. To compare more than two normally distributed sample groups, one-way ANOVA followed by post-hoc Tukey’s multiple comparisons test was used. p<0.05 was considered significant. Statistical analysis was carried out using Prism v9 (GraphPad Software).

### Modelling and MD Simulations of Kv7.2 Channels with a Starting Closed AG

To model the Kv7.2 pore with a closed AG, we used the cryo-EM structure of the human channel (hKv7.2, PDB ID: 7CR0, apo-state).^49^ This configuration has a conductive SF, as confirmed by the cross-distances between G279 Cα atoms of pairs of diagonally opposed monomers across the pore (distance d1= 8.03 Å). Moreover, the AG is characterized by short distances between Cα atoms of facing G313 (distance d2= 11.67 Å) and A317 (distance d3= 13.69 Å) residues.

Two other relevant closed-AG constriction points are identified by distances between S314 Oγ atoms (d4=4.00 Å) and L318 Cδ atoms (d5=7.14 Å). All five distances are illustrated in **Supplementary Fig. 4**. The segment of each subunit defining the TM pore domain (residues G215 to E330) was retained for MD simulations, while the VSDs were removed to reduce the size of the system. In light of DEE-related mutations affecting the D282 residue located close to the selectivity filter,^34,47,48^ the WT structure was simulated with either a charged or a protonated neutral aspartic acid at this site.

The CHARMM-GUI Membrane Builder server^64^ was used to prepare all necessary files for the simulations. Each of the five Kv7.2 structures (WT Kv7.2 with either charged or neutral D282; variants A317T, L318V and G313S) was inserted in a 1-palmitoyl-2-oleoyl-sn-glycero-3-phosphocholine (POPC) membrane and solvated with explicit water molecules. The total charge was neutralized with a 150 mM KCl solution, obtaining around 155,000 atoms in total. We used the NAMD software^65^ and the CHARMM36^66–68^ force field for proteins and lipids and using an integration time step Δt=2 fs. Before production, as already done in our previous works,^69,70^ all systems were relaxed with a ∼ 30 ns equilibration extending the CHARMM-GUI equilibration protocol, allowing proper hydration of solvent-exposed regions of the pore cavity. Then, five statistically independent replicates of each system were simulated for 500 ns each in the NPT ensemble at 1 atm and 310 K, respectively.

To investigate the effect of the membrane lipid composition, we performed additional MD simulations of the Kv7.2 A317T channel embedded in membranes of different lipid species (**Supplementary Table 1**), such as 1-palmitoyl-2-oleoyl-sn-glycero-3-phosphoethanolamine (POPE), 1-Palmitoyl-2-oleoyl-sn-glycero-3-(phospho-rac-(1-glycerol)) (POPG), 1,2-Dimyristoyl-sn-glycero-3-phosphocholine (DMPC), and a mixture POPC-Cholesterol and PIP_2_. To speed up calculations, the hydrogen mass repartitioning (HMR) method with an integration time step of 4 fs was used in these runs.^71–73^ Before production, each system was relaxed beyond the default CHARMM-GUI equilibration procedure with additional ∼70 ns in the NPT ensemble. Additional details about the MD set-up and accompanying results can be found in the **Supplementary Information**.

### Modeling and MD Simulations of the Kv7.2 Channels with a Starting Open Activation Gate

When we started this project, no experimental structure of Kv7.2 with an open IG was available. We thus relied on homology modeling using the open human Kv7.1 (hKv7.1, PDB ID: 6V01) as template.^38^ Our model comprises only the pore TM domain (residues G215 to E330), and was constructed in two steps following previously-described procedures.^69,70,74^ First, we modeled each of the four TM segments separately, using the structure prediction method included in the SWISS MODEL server.^75^ Then, the Kv7.2 pore domain was assembled by structurally aligning the four subunits to the template in a clockwise order viewed from the extracellular side, using UCSF Chimera.^76^ After the standard CHARMM-GUI relaxation, an additional 30 ns-long equilibration was performed with positional restraints on the protein and on the membrane atoms. The equilibrated channel has a conductive SF (d1= 7.97 Å) and a partially open AG (d2= 13.55 Å; d3= 19.62 Å). At this point, to avoid the sudden occlusion of the AG, a specific additional equilibration procedure was designed: three independent replicates of the system were further simulated for 500 ns, each by applying a set of harmonic restraints on the atomic distances between pairs of Cα atoms of residues G313 and A317 from diagonally opposed subunits, and then for 100 ns completely unrestrained. At the end of these simulations, the gate was stably open, and the final conformations were used as starting structures for five 500 ns-long MD simulations of each of the channels investigated.

### Calculations of the channel pore radius

The pore radius of each structure was calculated using the HOLE^77,78^ program. For replicated trajectories, average and standard deviations values were obtained from aggregating the five 500 ns-long runs and taking a snapshot every 50 ns. The structures were aligned to the starting conformation using the backbone atoms and excluding the terminal residues from 325 to 330 in each chain. The first 150 ns of each closed system were removed from the analysis to allow the mutated systems time to reach a stable conformation. For all the HOLE calculations, AMBER van der Waals radii were adopted with a cutoff of 6 Å.

### Hydration analysis

The hydration state of the different Kv7.2 pores was determined by calculating the number of water molecules along the pore axis during MD simulations. The pore axis, aligned along the z-axis, was divided into 20 intervals of 2.5 Å to map an interval of 50 Å comprising the whole cavity. In each interval, the average number of water molecules and standard deviations were calculated using VMD and Tcl scripting.^79^ As for the pore radii calculations, the first 150 ns of each closed IG system were discarded. For replicated trajectories, the mean and standard deviation were extracted from the concatenated runs.

### Hydrogen bond analysis

The hydrogen bond (HB) analysis was performed using VMD. HBs were defined with a cutoff distance between the heavy atoms covalently bound to a hydrogen atom of 3.3 Å, and an angle defined by the heavy atoms and the central hydrogen atom comprised between 130° and 230°.

### Molecular graphics

Molecular graphics and further analysis were performed with UCSF ChimeraX^80–82^ and Visual Molecular Dynamics (VMD).

## AUTHOR CONTRIBUTION

Conceptualization: MN, GA, LM, SW, IES, FM, MT; Methodology: MN, GA, LM, FM; Data curation: GA, AB, AR, TGAC, IES; Investigation: MN, GA, AB, AR, MVS, VB, TGAC, IES, LM, FM,; Visualization: MN, GA, AB, AR, MVS, VB, FM; Funding acquisition: GA, LM, FB, FM, MT; Writing – original draft: MN, GA, FM, MT; Writing - review & editing: all authors.

## ACKNOWLEDGMENTS

We thank Alessia Vignolo, Mattia Pini and Sergio Decherchi for the kind assistance at the Italian Institute of Technology (IIT) computing center, and Michael Pusch at Biophysics Institute at the National Research Council (CNR) in Genova (IT) for sharing noise analysis softwares. We are grateful to Diego Moruzzo for useful help and technical assistance. Computing time allocations were granted by the CINECA supercomputing center under the ISCRA initiative. We also gratefully acknowledge the High Performance Computing (HPC) infrastructure and the Support Team at Fondazione IIT.

This work was supported by the Italian Ministry for University and Research (MUR) with PRIN2020 (Project 2020XBFEMS to LM), PRIN2022 (Project 2022M3KJ4N to MT), PRIN2022PNRR (project P2022FJXY5 to MVS and FM; project P2022ZANRF to MT), the MNESYS project “A multiscale integrated approach to the study of the nervous system in health and disease” of the Next Generation EU National Recovery and Resilience Plan (to MT), by the Italian Ministry of Health (MoH) with Ricerca Finalizzata (RF) Projects: RF-2019-12370491 to MT, and PNRR-MR1-2022-12376528 to MVS, FB, and MT, by the FWO (grants 1861419N and G041821N to SW), and by the European Joint Programme on Rare Disease JTC 2020 (TreatKCNQ, to SW and MT).

The DDD study presents independent research commissioned by the Health Innovation Challenge Fund [grant number HICF-1009-003]. This study makes use of DECIPHER (http://www.deciphergenomics.org), which is funded by Wellcome [grant number WT223718/Z/21/Z]. See Ref. 83 or www.ddduk.org/access.html for full acknowledgement.

## ABBREVIATIONS

CHOL: cholesterol
CD: cross-distance
CHO: Chinese Hamster Ovary
Cryo-EM: cryogenic electron microscopy
DEE: developmental and epileptic encephalopathy
DMPC: 1,2-Dimyristoyl-sn-glycero-3-phosphocholine
ESES: 
GoF: gain-of-function
HB: hydrogen bond
IG: inner gate
Kv: voltage-gated potassium channel
LoF: loss-of-function
MD: molecular dynamics
NDD: neurodevelopmental disorder
PBS: phosphate-buffered saline
PAG: pore activation gate
PD: pore domain
PIP2: phosphatidylinositol 4,5-bisphosphate
POPC: 1-palmitoyl-2-oleoyl-sn-glycero-3-phosphocholine
POPE: 1-palmitoyl-2-oleoyl-sn-glycero-3-phosphoethanolamine
POPG: 1-Palmitoyl-2-oleoyl-sn-glycero-3-(phospho-rac-(1-glycerol))
RMSD: root mean square deviation
SDS-PAGE: sodium dodecylsulfate polyacrylamide gel electrophoresis
SF: selectivity filter
SLFNE: self-limited familial neonatal epilepsy
TM: transmembrane
VSD: voltage-sensing domain
WT: wild type

## REFERENCES

1. Weckhuysen, S. et al. KCNQ2 encephalopathy: Emerging phenotype of a neonatal epileptic encephalopathy. Annals of Neurology 71, 15–25 (2012).

2. Nappi, P. et al. Epileptic channelopathies caused by neuronal Kv7 (KCNQ) channel dysfunction. Pflugers Arch 472, 881–898 (2020).

3. Miceli, F., Soldovieri, M. V., Weckhuysen, S., Cooper, E. & Taglialatela, M. KCNQ2-Related Disorders. in GeneReviews® (eds. Adam, M. P. et al.) (University of Washington, Seattle, Seattle (WA), 1993).

4. Miceli, F., Soldovieri, M. V., Weckhuysen, S., Cooper, E. C. & Taglialatela, M. KCNQ3-Related Disorders. in GeneReviews® (eds. Adam, M. P. et al.) (University of Washington, Seattle, Seattle (WA), 1993).

5. Zuberi, S. M. et al. ILAE classification and definition of epilepsy syndromes with onset in neonates and infants: Position statement by the ILAE Task Force on Nosology and Definitions. Epilepsia 63, 1349– 1397 (2022).

6. Scheffer, I. E. et al. ILAE classification of the epilepsies: Position paper of the ILAE Commission for Classification and Terminology. Epilepsia 58, 512–521 (2017).

7. Symonds, J. D. et al. Incidence and phenotypes of childhood-onset genetic epilepsies: a prospective population-based national cohort. Brain 142, 2303–2318 (2019).

8. Mary, L. et al. Pathogenic variants in KCNQ2 cause intellectual deficiency without epilepsy: Broadening the phenotypic spectrum of a potassium channelopathy. American Journal of Medical Genetics Part A 185, 1803–1815 (2021).

9. Barrese, V., Stott, J. B. & Greenwood, I. A. KCNQ-Encoded Potassium Channels as Therapeutic Targets. Annual Review of Pharmacology and Toxicology 58, 625–648 (2018).

10. Soldovieri, M. V., Miceli, F. & Taglialatela, M. Driving With No Brakes: Molecular Pathophysiology of Kv7 Potassium Channels. Physiology 26, 365–376 (2011).

11. Huang, Y., Ma, D., Yang, Z., Zhao, Y. & Guo, J. Voltage-gated potassium channels KCNQs: Structures, mechanisms, and modulations. Biochemical and Biophysical Research Communications 689, 149218 (2023).

12. Gu, R.X. & de Groot, B.L. Central cavity dehydration as a gating mechanism of potassium channels. Nat Commun. 14,: 2178 (2023).

13. Zaydman, M. A. & Cui, J. PIP2 regulation of KCNQ channels: biophysical and molecular mechanisms for lipid modulation of voltage-dependent gating. Front Physiol 5, 195 (2014).

14. Zheng, Y. et al. Structural insights into the lipid and ligand regulation of a human neuronal KCNQ channel. Neuron 110, 237–247.e4 (2022).

15. Ma, D. et al. Ligand activation mechanisms of human KCNQ2 channel. Nat Commun 14, 6632 (2023).

16. Rosa, F. et al. Electrophysiological signatures of a developmental delay in a stem cell model of KCNQ2 developmental and epileptic encephalopathy. 2024.03.13.584717 Preprint at 10.1101/2024.03.13.584717 (2024).

17. Miceli, F. et al. Genotype-phenotype correlations in neonatal epilepsies caused by mutations in the voltage sensor of K(v)7.2 potassium channel subunits. Proc Natl Acad Sci U S A 110, 4386–4391 (2013).

18. Orhan, G. et al. Dominant-negative effects of KCNQ2 mutations are associated with epileptic encephalopathy. Ann Neurol 75, 382–394 (2014).

19. Brun, L., Viemari, J.-C. & Villard, L. Mouse models of Kcnq2 dysfunction. Epilepsia 63, 2813–2826 (2022).

20. Millichap, J. J. et al. Infantile spasms and encephalopathy without preceding neonatal seizures caused by KCNQ2 R198Q, a gain-of-function variant. Epilepsia 58, e10–e15 (2017).

21. Bayat, A. et al. Phenotypic and Functional Assessment of Two Novel KCNQ2 Gain-of-Function Variants Y141N and G239S and Effects of Amitriptyline Treatment. https://www.researchsquare.com/article/rs-2710358/v1 (2023) doi:10.21203/rs.3.rs-2710358/v1.

22. Miceli, F. et al. KCNQ2 R144 variants cause neurodevelopmental disability with language impairment and autistic features without neonatal seizures through a gain-of-function mechanism. eBioMedicine 81, 104130 (2022).

23. Mulkey, S. B. et al. Neonatal nonepileptic myoclonus is a prominent clinical feature of KCNQ2 gain-of-function variants R201C and R201H. Epilepsia 58, 436–445 (2017).

24. Sands, T. T. et al. Autism and developmental disability caused by KCNQ3 gain-of-function variants. Ann Neurol 86, 181–192 (2019).

25. Niday, Z. & Tzingounis, A. V. Potassium Channel Gain of Function in Epilepsy: An Unresolved Paradox. Neuroscientist 24, 368–380 (2018).

26. Varghese, N. et al. KCNQ2/3 Gain-of-Function Variants and Cell Excitability: Differential Effects in CA1 versus L2/3 Pyramidal Neurons. J. Neurosci. 43, 6479–6494 (2023).

27. Devaux, J. et al. A Kv7.2 mutation associated with early onset epileptic encephalopathy with suppression-burst enhances Kv7/M channel activity. Epilepsia 57, e87–93 (2016).

28. Miceli, F. et al. Early-onset epileptic encephalopathy caused by gain-of-function mutations in the voltage sensor of Kv7.2 and Kv7.3 potassium channel subunits. J Neurosci 35, 3782–3793 (2015).

29. Edmond, M. A., Hinojo-Perez, A., Wu, X., Perez Rodriguez, M. E. & Barro-Soria, R. Distinctive mechanisms of epilepsy-causing mutants discovered by measuring S4 movement in KCNQ2 channels. eLife 11, e77030 (2022).

30. Barro-Soria, R. Epilepsy-associated mutations in the voltage sensor of KCNQ3 affect voltage dependence of channel opening. Journal of General Physiology 151, 247–257 (2018).

31. Masnada, S. et al. Clinical spectrum and genotype–phenotype associations of KCNA2-related encephalopathies. Brain 140, 2337–2354 (2017).

32. Brunklaus A, et al. Gene variant effects across sodium channelopathies predict function and guide precision therapy. Brain 145, 4275–4286 (2022).

33. Gong, P., Xue, J., Jiao, X., Zhang, Y. & Yang, Z. Genetic Etiologies in Developmental and/or Epileptic Encephalopathy With Electrical Status Epilepticus During Sleep: Cohort Study. Front. Genet. 12, 607965 (2021).

34. Deciphering Developmental Disorders Study. Prevalence and architecture of de novo mutations in developmental disorders. Nature 542, 433–438 (2017).

35. Specchio, N. et al. International League Against Epilepsy classification and definition of epilepsy syndromes with onset in childhood: Position paper by the ILAE Task Force on Nosology and Definitions. Epilepsia 63, 1398–1442 (2022).

36. Richards, S. et al. Standards and guidelines for the interpretation of sequence variants: a joint consensus recommendation of the American College of Medical Genetics and Genomics and the Association for Molecular Pathology. Genetics in Medicine 17, 405–424 (2015).

37. Li, X. et al. Molecular basis for ligand activation of the human KCNQ2 channel. Cell Res 31, 52–61 (2021).

38. Sun, J. & MacKinnon, R. Structural Basis of Human KCNQ1 Modulation and Gating. Cell 180, 340–347.e9 (2020).

39. Hadley, J. K. et al. Differential tetraethylammonium sensitivity of KCNQ1–4 potassium channels. British Journal of Pharmacology 129, 413–415 (2000).

40. Alvarez, O., Gonzalez, C. & Latorre, R. Counting channels: a tutorial guide on ion channel fluctuation analysis. Advances in Physiology Education 26, 327–341 (2002).

41. Kim, R. Y., Pless, S. A. & Kurata, H. T. PIP2 mediates functional coupling and pharmacology of neuronal KCNQ channels. Proceedings of the National Academy of Sciences 114, E9702–E9711 (2017).

42. Pant, S. et al. PIP2-dependent coupling of voltage sensor and pore domains in Kv7.2 channel. Commun Biol 4, 1189 (2021).

43. Falkenburger, B. H., Jensen, J. B. & Hille, B. Kinetics of PIP2 metabolism and KCNQ2/3 channel regulation studied with a voltage-sensitive phosphatase in living cells. J Gen Physiol 135, 99–114 (2010).

44. Soldovieri, M. V. et al. Early-onset epileptic encephalopathy caused by a reduced sensitivity of Kv7.2 potassium channels to phosphatidylinositol 4,5-bisphosphate. Sci Rep 6, 38167 (2016).

45. Hossain, M. I. et al. Enzyme domain affects the movement of the voltage sensor in ascidian and zebrafish voltage-sensing phosphatases. J Biol Chem 283, 18248–18259 (2008).

46. Nappi, M. et al. Gain of function due to increased opening probability by two KCNQ5 pore variants causing developmental and epileptic encephalopathy. Proceedings of the National Academy of Sciences of the United States of America 119, (2022).

47. Millichap, John J., et al. KCNQ2 encephalopathy: Features, mutational hot spots, and ezogabine treatment of 11 patients. Neurology: Genetics 2, e96 (2016).

48. Olson, H.E., et al. Genetics and genotype–phenotype correlations in early onset epileptic encephalopathy with burst suppression. Annals of Neurology 81, 419–429 (2017).

49. Li, X., et al. Molecular basis for ligand activation of the human KCNQ2 channel. Cell Res 31, 52–61 (2021).

50. Zhang, S., et al. A small-molecule activation mechanism that directly opens the KCNQ2 channel. Nat Chem Biol 1–10 (2024).

51. Miceli, F. et al. A novel KCNQ3 mutation in familial epilepsy with focal seizures and intellectual disability. Epilepsia 56, e15–e20 (2015).

52. Ambrosino, P. et al. Kv7.3 Compound Heterozygous Variants in Early Onset Encephalopathy Reveal Additive Contribution of C-Terminal Residues to PIP_2_-Dependent K^+^ Channel Gating. Mol Neurobiol 55, 7009–7024 (2018).

53. Sun, J. & MacKinnon, R. Cryo-EM Structure of a KCNQ1/CaM Complex Reveals Insights into Congenital Long QT Syndrome. Cell 169, 1042–1050.e9 (2017).

54. Mandala, V. S. & MacKinnon, R. The membrane electric field regulates the PIP2-binding site to gate the KCNQ1 channel. Proceedings of the National Academy of Sciences 120, e2301985120 (2023).

55. Willegems, K. et al. Structural and electrophysiological basis for the modulation of KCNQ1 channel currents by ML277. Nat Commun 13, 3760 (2022).

56. Ma, D. et al. Structural mechanisms for the activation of human cardiac KCNQ1 channel by electro-mechanical coupling enhancers. Proceedings of the National Academy of Sciences 119, e2207067119 (2022).

57. Li, T. et al. Structural Basis for the Modulation of Human KCNQ4 by Small-Molecule Drugs. Molecular Cell 81, 25–37.e4 (2021).

58. Zaika, O., Hernandez, C. C., Bal, M., Tolstykh, G. P. & Shapiro, M. S. Determinants within the Turret and Pore-Loop Domains of KCNQ3 K+ Channels Governing Functional Activity. Biophysical Journal 95, 5121–5137 (2008).

59. Zhang, H. et al. PIP(2) activates KCNQ channels, and its hydrolysis underlies receptor-mediated inhibition of M currents. Neuron 37, 963–975 (2003).

60. Mattmann, M. E. et al. Identification of (R)-N-(4-(4-methoxyphenyl)thiazol-2-yl)-1-tosylpiperidine-2-carboxamide, ML277, as a novel, potent and selective K(v)7.1 (KCNQ1) potassium channel activator. Bioorg Med Chem Lett 22, 5936–5941 (2012).

61. Brennan, S., et al. Slowly activating voltage-gated potassium current potentiation by ML277 is a novel cardioprotective intervention. PNAS Nexus 2, pgad156 (2023).

62. Soldovieri, M. V. et al. Atypical gating of M-type potassium channels conferred by mutations in uncharged residues in the S4 region of KCNQ2 causing benign familial neonatal convulsions. J Neurosci 27, 4919–4928 (2007).

63. Prole, D. L. & Marrion, N. V. Ionic Permeation and Conduction Properties of Neuronal KCNQ2/KCNQ3 Potassium Channels. Biophysical Journal 86, 1454–1469 (2004).

64. Jo, S., Kim, T., & Im, W. Automated builder and database of protein/membrane complexes for molecular dynamics simulations. PloS one 2, e880 (2007).

65. Phillips, J. C. et al. Scalable molecular dynamics with NAMD. J Comput Chem 26, 1781–1802 (2005).

66. Best, R. B. et al. Optimization of the Additive CHARMM All-Atom Protein Force Field Targeting Improved Sampling of the Backbone -, ψ and Side-Chain χ1 and χ2 Dihedral Angles. J. Chem. Theory Comput. 8, 3257–3273 (2012).

67. Huang, J. & MacKerell, A. D. CHARMM36 all-atom additive protein force field: validation based on comparison to NMR data. J Comput Chem 34, 2135–2145 (2013).

68. Klauda, J. B. et al. Update of the CHARMM All-Atom Additive Force Field for Lipids: Validation on Six Lipid Types. J. Phys. Chem. B 114, 7830–7843 (2010).

69. Alberini, G., Benfenati, F. & Maragliano, L. Structural Mechanism of ω-Currents in a Mutated Kv7.2 Voltage Sensor Domain from Molecular Dynamics Simulations. J. Chem. Inf. Model. 61, 1354–1367 (2021).

70. Alberini, G. et al. Molecular Dynamics Simulations of Ion Permeation in Human Voltage-Gated Sodium Channels. J. Chem. Theory Comput. 19, 2953–2972 (2023).

71. Feenstra, K. A., Hess, B. & Berendsen, H. J. C. Improving efficiency of large time-scale molecular dynamics simulations of hydrogen-rich systems. Journal of Computational Chemistry 20, 786–798 (1999).

72. Hopkins, C. W., Le Grand, S., Walker, R. C. & Roitberg, A. E. Long-Time-Step Molecular Dynamics through Hydrogen Mass Repartitioning. J. Chem. Theory Comput. 11, 1864–1874 (2015).

73. Balusek, C. et al. Accelerating Membrane Simulations with Hydrogen Mass Repartitioning. J Chem Theory Comput 15, 4673–4686 (2019).

74. Yang, Y. et al. Structural modelling and mutant cycle analysis predict pharmacoresponsiveness of a Nav1.7 mutant channel. Nat Commun 3, 1186 (2012).

75. Waterhouse, A. et al. SWISS-MODEL: homology modelling of protein structures and complexes. Nucleic Acids Research 46, W296–W303 (2018).

76. Pettersen, E. F. et al. UCSF Chimera—A visualization system for exploratory research and analysis. Journal of Computational Chemistry 25, 1605–1612 (2004).

77. Smart, O. S., Neduvelil, J. G., Wang, X., Wallace, B. A. & Sansom, M. S. P. HOLE: A program for the analysis of the pore dimensions of ion channel structural models. Journal of Molecular Graphics 14, 354–360 (1996).

78. Smart, O. S., Breed, J., Smith, G. R. & Sansom, M. S. A novel method for structure-based prediction of ion channel conductance properties. Biophysical Journal 72, 1109 (1997).

79. Humphrey, W., Dalke, A. & Schulten, K. VMD: Visual molecular dynamics. Journal of Molecular Graphics 14, 33–38 (1996).

80. Goddard, T. D. et al. UCSF ChimeraX: Meeting modern challenges in visualization and analysis. Protein Sci 27, 14–25 (2018).

81. Meng, E. C. et al. UCSF ChimeraX: Tools for structure building and analysis. Protein Sci 32, e4792 (2023).

82. Pettersen, E. F. et al. UCSF ChimeraX: Structure visualization for researchers, educators, and developers. Protein Sci 30, 70–82 (2021).

83. Deciphering Developmental Disorders Study. Large-scale discovery of novel genetic causes of developmental disorders. Nature 519, 223-228 (2015).

