## Supplementary Figures and Methods for "CONSTITUTIVE OPENING OF THE Kv7.2 PORE ACTIVATION GATE CAUSES *KCNQ2*-DEVELOPMENTAL ENCEPHALOPATHY"

**CONTENT:**

**Supplementary Figs. 1-3: Supplementary data for functional analysis**
**Supplementary Methods for Molecular Dynamics**
**Supplementary Table 1: Summary of the MD simulations**
**Supplementary Figs. 4-24: Supplementary for Molecular Dynamics**
**Bibliography**

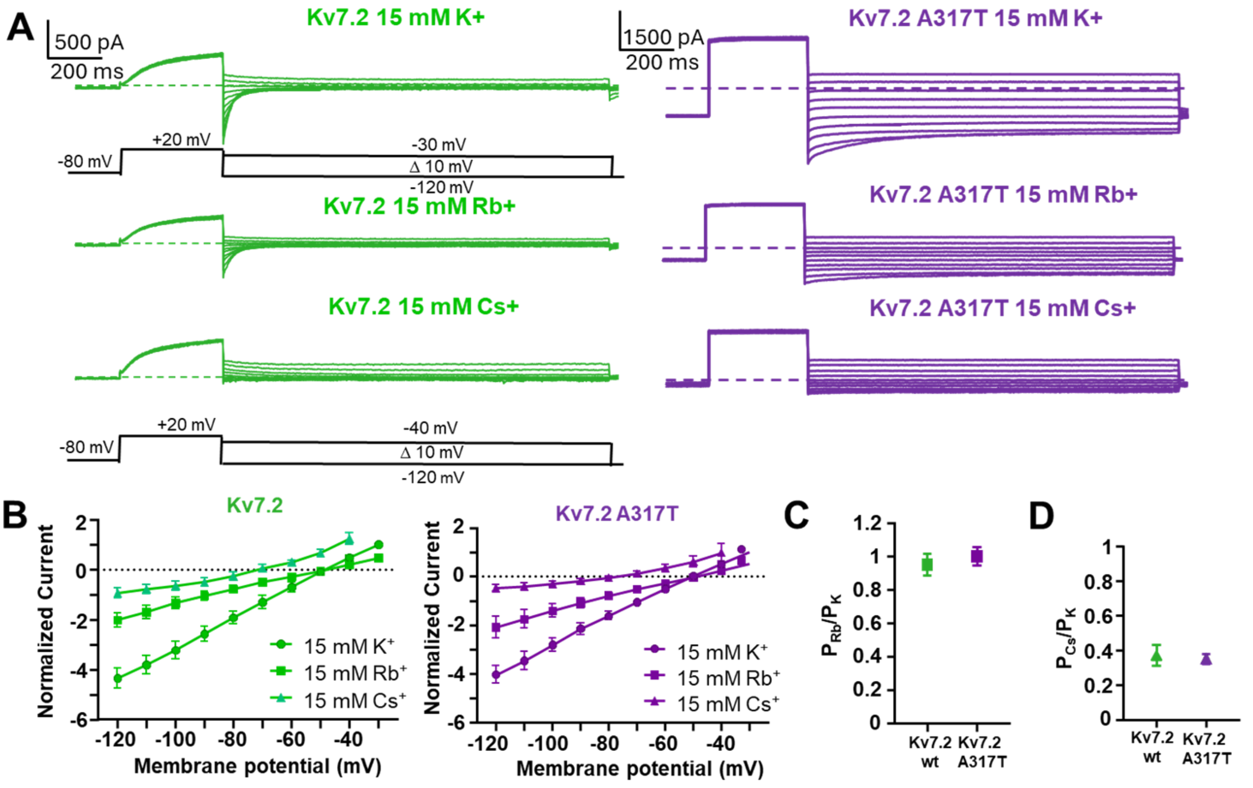


**E**

| | | $Rb^+$ | $Cs^+$ |
|---|---|---|---|
| $P_X/P_K$ | **Kv7.2** | 0.95 ± 0.07 | 0.37 ± 0.06 |
| | **Kv7.2 A317T** | 1.00 ± 0.05 | 0.35 ± 0.10 |

**Supplementary Fig. 1. Permeation properties of Kv7.2 and Kv7.2 A317T channels.**
**A.** Representative traces from wild type Kv7.2 (green) and Kv7.2 A317T (purple) in the presence of extracellular potassium (upper traces), rubidium (middle traces) and caesium (bottom traces) are reported. To measure reversal potentials in the presence of $K^+$, $Rb^+$, and $Cs^+$, membrane potential were clamped as reported in the corresponding voltage protocols. **B.** Current-voltage relationships for Kv7.2 and Kv7.2 A317T currents in the presence of 15 mM external $K^+$, $Rb^+$, and $Cs^+$+. All the current values were normalized to the values obtained at +30 mV in 15 mM K+ in the same cell. N=6-8. Reversal potential values (mean ± SEM, N=6-8), expressed in mV, were -47.41±0.94 in 15 mM $K^+$, 49.17±2.02 in 15 mM $Rb^+$, and -73.38±4.60 in 15 mM $Cs^+$ for WT Kv7.2; -50.03±1.22 in 15 mM $K^+$, 50.18±1.47 in 15 mM $Rb^+$, and -79.96±6.17 in 15 mM $Cs^+$ for Kv7.2 A317T. **C** and **D** show cations permeability rations ($P_{Rb}/P_K$ and $P_{Cs}/P_K$ respectively), calculated as indicated in the Method section. The ratio between the permeability of potassium and the permeability of rubidium or caesium was calculated on the same cell. In **E**, permeability ratios numerical values are reported.

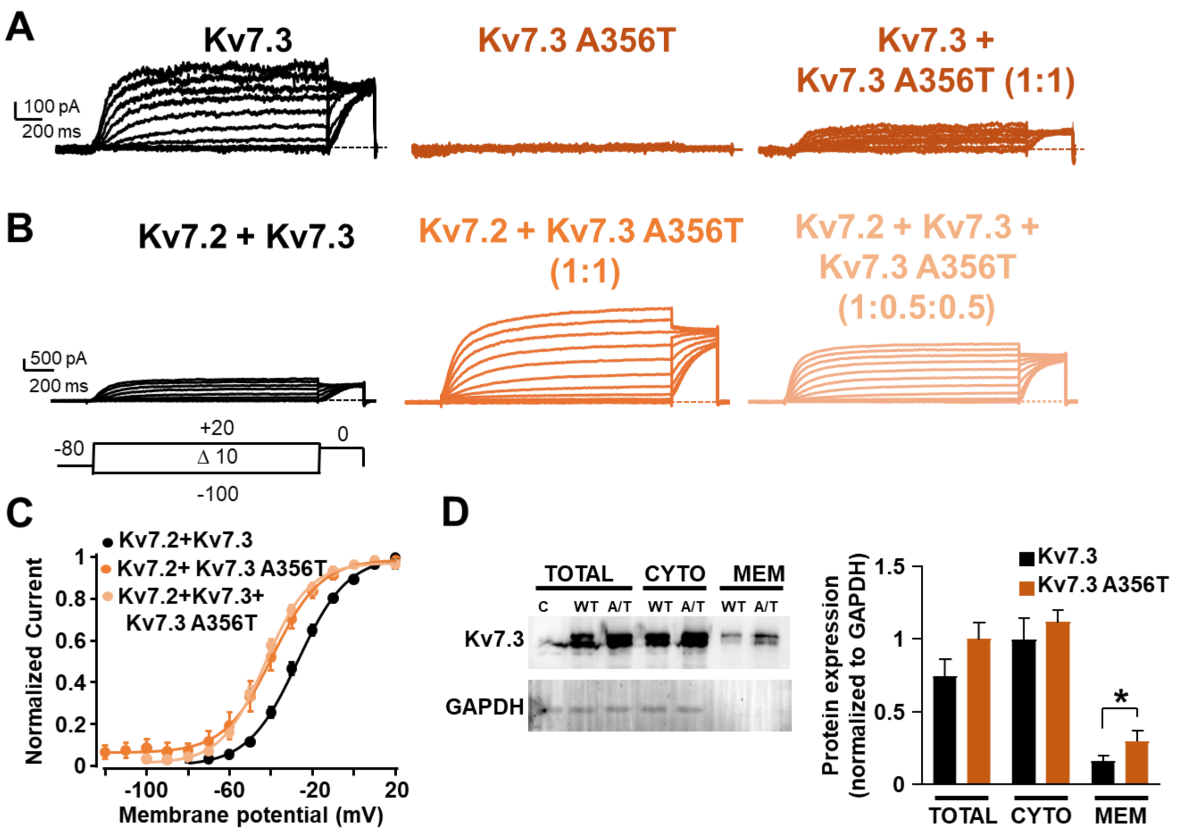


**Supplementary Fig. 2. Functional characterization of Kv7.3 A356T currents in both homomeric and heteromeric assembly. A.** Currents from Kv7.3, Kv7.3 A356T and Kv7.3 + Kv7.3 A356T (1:1 cDNA ratio). **B.** Currents from Kv7.2 + Kv7.3, Kv7.2 + Kv7.3 A356T, and Kv7.2 + Kv7.3 + Kv7.3 A356T channels. cDNA ratios are indicate in parenthesis. **C.** Conductance/voltage curves for the indicated channels; continuous lines represent Boltzmann fits of the experimental data. **D.** Representative Western-blot experiments of proteins from total, cytosol (CYTO) or plasma membrane (MEM) fractions from CHO cells transfected with pcDNA3.1 (empty vector, C) Kv7.3 or Kv7.3 A356T (A/T) subunits. Lysates were incubated with anti-Kv7.3 antibody (top image) or anti-GAPDH antibody (lower image), as indicated. The right panel reports the densitometric quantification of the Kv7.3 band intensity (normalized to that corresponding to GAPDH) for the two experimental groups. Data are expressed as Mean±S.E.M. (n=4).

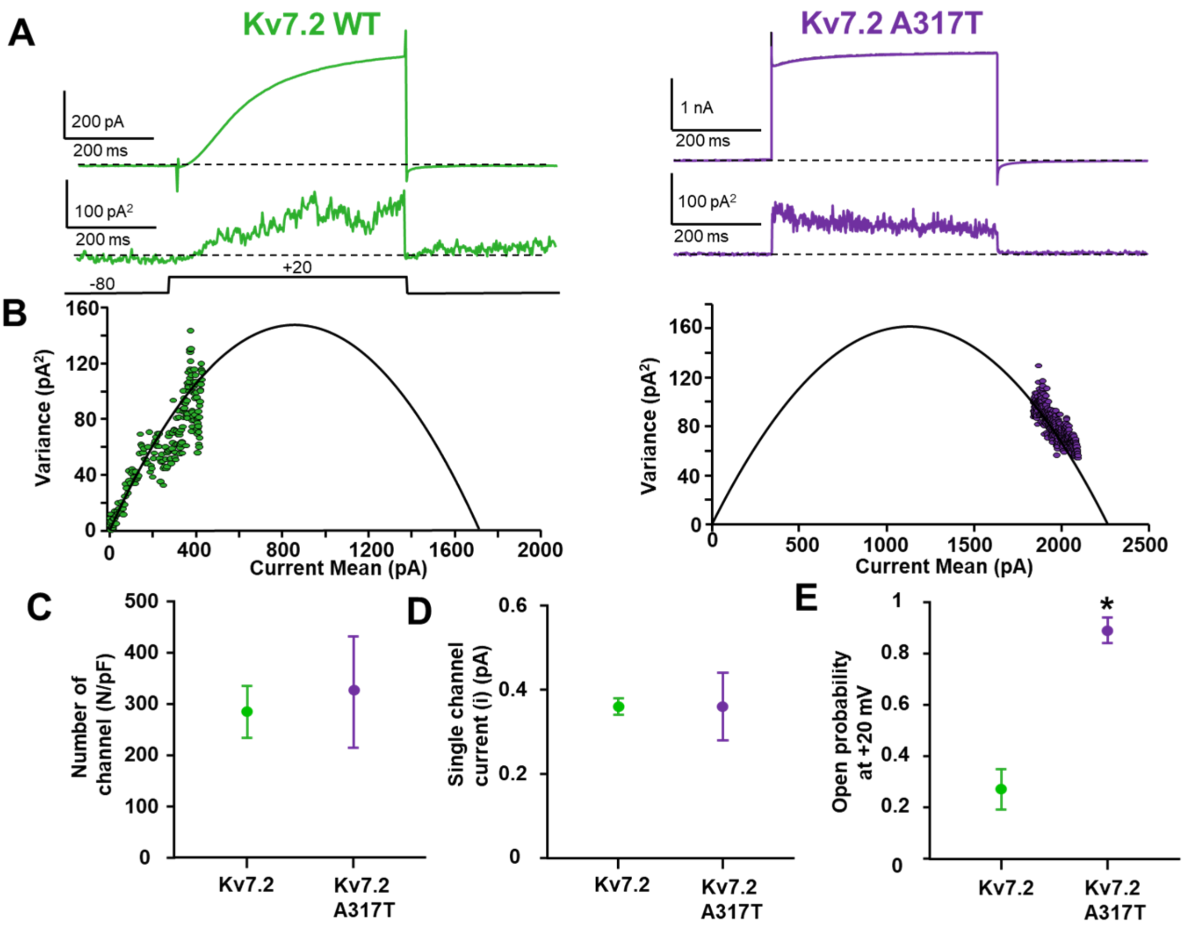


**Supplementary Fig. 3. Non-stationary noise analysis of the K$^+$ currents from Kv7.2A317T channels**. **A.** Representative average current responses to 100 pulses at +20 mV (top traces) and of the respective variance (bottom traces) for the indicated channels. **B**. Variance versus current mean plot derived from Kv7.2 (green) and Kv7.2 A317T (purple). The continuous lines in the variance/mean plots are parabolic fits of the experimental data to equation 2 in Methods. Panels **C**–**E** depict the quantification of the number of channels divided by capacitance (**C**), of the single-channel current (**D**), and of the opening probability at +20 mV (**E**) for the indicated channels. The asterisk highlights a value significantly different (*$p < 0.05$) versus controls (Kv7.2).

### SUPPLEMENTARY METHODS FOR MOLECULAR DYNAMICS

**Modelling and simulations of Kv7.2 channels with a starting closed activation gate.**

The CHARMM-GUI Membrane Builder server[1] was used to prepare all necessary files for the simulations. Before MD simulations, all the systems were oriented within the membrane using the OPM-PPM[2] server. The NAMD software[3] and the CHARMM36[4-6] force field for proteins and lipids, along with the CHARMM-modified TIP3P model for water molecules[7]. CHARMM-compatible ionic parameters with NBFIX corrections were employed[8-10]. Tetragonal periodic boundary conditions (PBCs) were applied to the simulation box to remove surface effects, and the Particle Mesh Ewald (PME) method was used to calculate long range electrostatic interactions[11]. Short-range electrostatic and van der Waals interactions were calculated with a 12 Å cutoff and by applying a smooth decaying function starting to take effect at 10 Å. The CHARMM force-based switching function was employed for van der Waals interactions via the vdwForceSwitching command.[12] The SHAKE algorithm[13] was used to constrain covalent bonds involving hydrogen atoms (except for water molecules, where SETTLE[14] was used), allowing for an integration time step $\Delta t$=2 fs. To ensure maximum accuracy, electrostatic and van der Waals interactions were computed at each simulation step.

We used the Nosé-Hoover Langevin piston method[15,16] and a Langevin thermostat to reproduce the NPT ensemble and maintain the pressure at 1 atm and the temperature at 310 K, respectively. Following the CHARMM-GUI input files, the oscillation period of the piston was set at 50 fs, and the damping time scale at 25 fs. The Langevin thermostat was set with a damping coefficient of 1 $ps^{-1}$.

For the additional membrane models simulations (**Supplementary Table 1**), each system was run for 500 ns. The NAMD software was used with the CHARMM36m[17]/CHARMM36 force field for protein and lipids, respectively. Long range electrostatic interactions were calculated using the Particle Mesh Ewald (PME) algorithm. Electrostatic and van der Waals interactions were calculated with a cutoff of 12 Å and the application of a smoothing decay starting to take effect at 10 Å.[18] Before production, each system was relaxed beyond the default CHARMM-GUI equilibration procedure with additional ∼70 ns in the NPT ensemble. The Nosé-Hoover Langevin piston method to maintain the pressure at 1 atm and a Langevin thermostat at 310K, above the phase transition temperature of each lipid (**Table S1**). The oscillation period of the piston was set at 300 fs, and the damping time scale at 150 fs[18]. The Langevin thermostat was employed with a damping coefficient of 1 $ps^{-1}$.

**Supplementary Table 1. Summary of the MD simulations.**

| System | IG | Starting conformation | Membrane | FF | Δt (fs) | Time (ns) |
|---|---|---|---|---|---|---|
| WT | Closed | cryo-EM | POPC | CHARMM36 | 2 | 500 x 5 |
| WT-D$^0$282 | Closed | cryo-EM | POPC | CHARMM36 | 2 | 500 x 5 |
| G313S | Closed | cryo-EM | POPC | CHARMM36 | 2 | 500 x 5 |
| A317T | Closed | cryo-EM | POPC | CHARMM36 | 2 | 500 x 5 |
| A317T | Closed | cryo-EM | POPE | CHARMM36m | 4 | 500 x 1 |
| A317T | Closed | cryo-EM | POPG | CHARMM36m | 4 | 500 x 1 |
| A317T | Closed | cryo-EM | POP-CHOL-PIP2 | CHARMM36m | 4 | 500 x 1 |
| A317T | Closed | cryo-EM | DMPC | CHARMM36m | 4 | 500 x 1 |
| L318V | Closed | cryo-EM | POPC | CHARMM36 | 2 | 500 x 5 |
| WT | Open | Hom. Mod. | POPC | CHARMM36 | 2 | 500 x 5 |
| G313S | Open | Hom. Mod. | POPC | CHARMM36 | 2 | 500 x 5 |
| A317T | Open | Hom. Mod. | POPC | CHARMM36 | 2 | 500 x 5 |
| L318V | Open | Hom. Mod. | POPC | CHARMM36 | 2 | 500 x 5 |

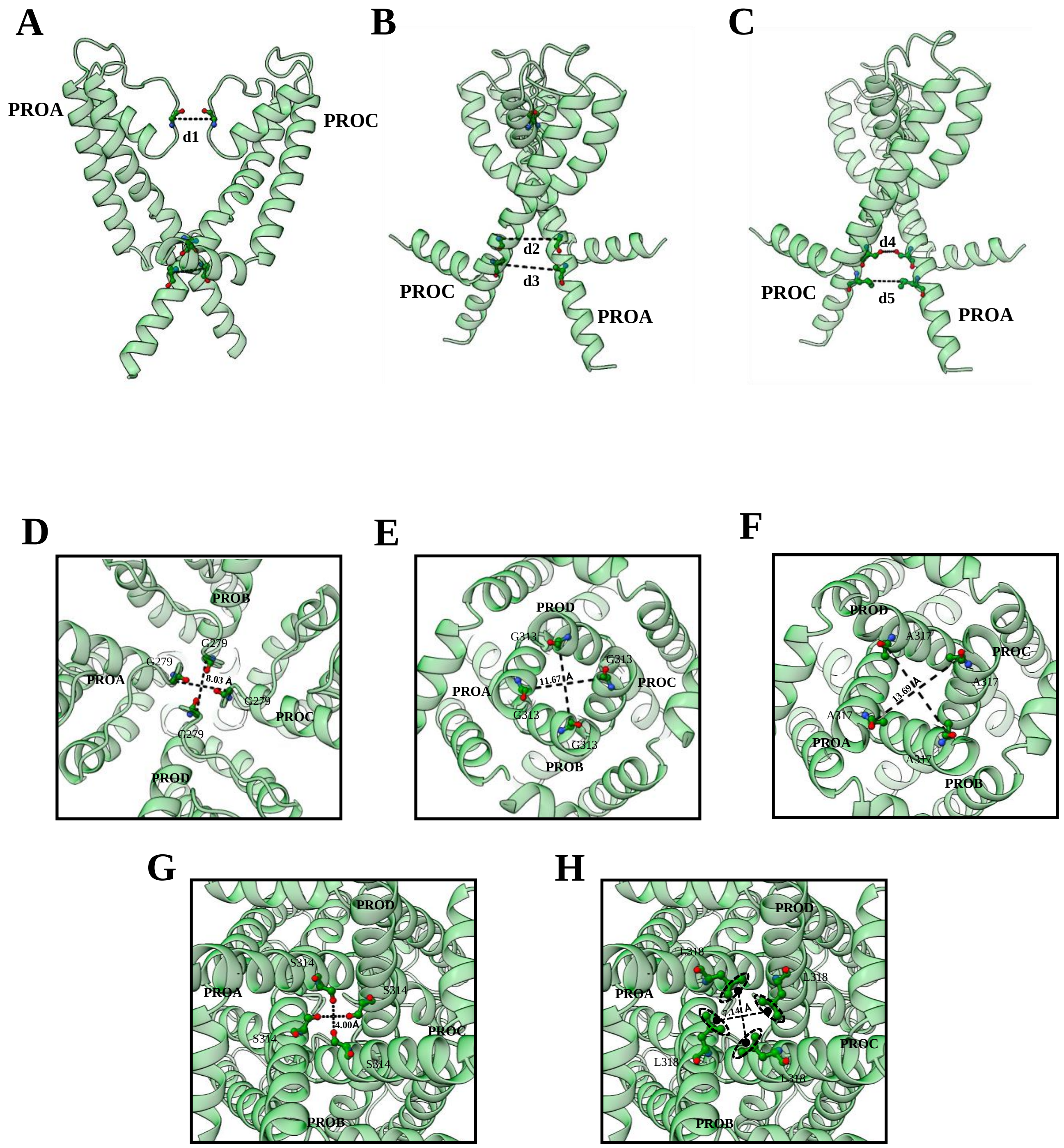


**Supplementary Fig. 4. Representation of cross-distances (CDs) d1 to d5 using the hKv7.2 cryo-EM structure (PDB ID: 7CR0). (A-C**) Lateral views from different angles, showing only two subunits. The residues employed to compute the CDs are shown as balls and sticks. **(D)** Distance d1, viewed from the extracellular side. **(E-H)** Distances d2 to d5, viewed from the cytosolic side.

**A**

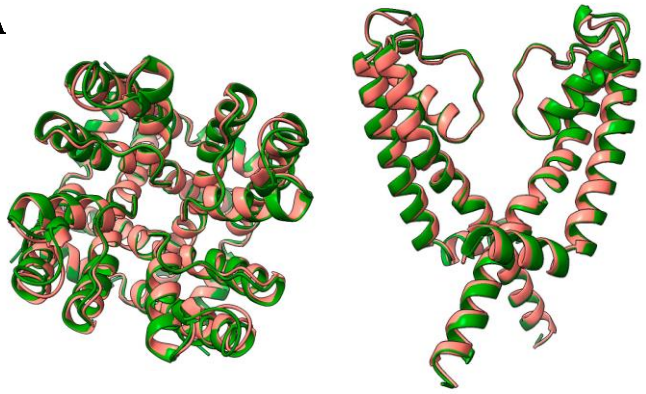

PDB ID: 8J01

| Distance | Average value | HM value |
| --- | --- | --- |
| d1(Cα$^{279}$) | 8.00 Å | 7.97 Å |
| d2(Cα$^{313}$) | 13.77 Å | 13.55 Å |
| d3(Cα$^{317}$) | 18.20 Å | 19.62 Å |

| | |
| --- | --- |
| **RMSD** | 0.889 Å |

**B**

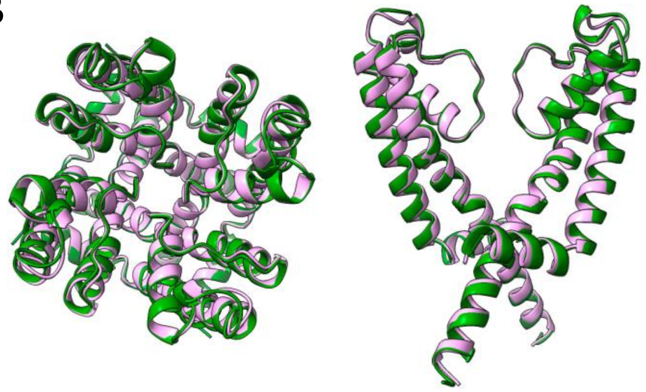

PDB ID: 8J02

| Distance | Average value |
| --- | --- |
| d1(Cα$^{279}$) | 7.95 Å |
| d2(Cα$^{313}$) | 13.70 Å |
| d3(Cα$^{317}$) | 17.72 Å |

| | |
| --- | --- |
| **RMSD** | 0.813 Å |

**C**

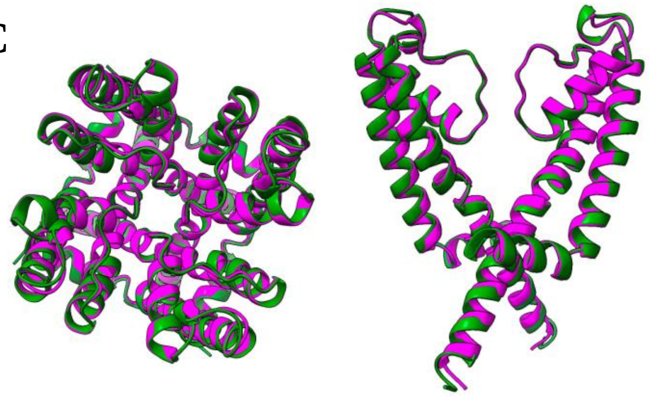

PDB ID: 8W4U

| Distance | Average value |
| --- | --- |
| d1(Cα$^{279}$) | 8.02 Å |
| d2(Cα$^{313}$) | 13.85 Å |
| d3(Cα$^{317}$) | 17.52 Å |

| | |
| --- | --- |
| **RMSD** | 0.846 Å |

**D**

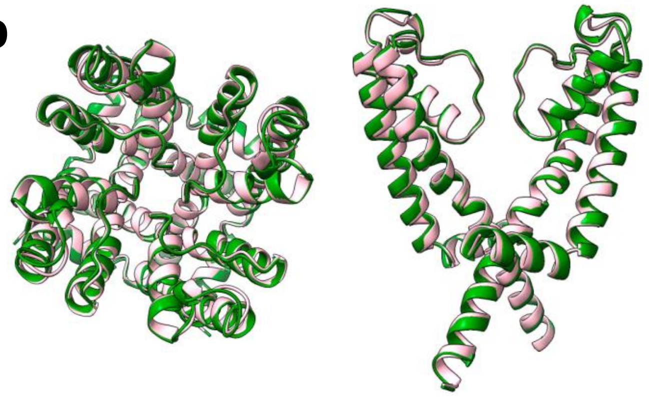

PDB ID: 8IJK

| Distance | Average value |
| --- | --- |
| d1(Cα$^{279}$) | 8.33 Å |
| d2(Cα$^{313}$) | 13.49 Å |
| d3(Cα$^{317}$) | 17.76 Å |

| | |
| --- | --- |
| **RMSD** | 0.331 Å |

**Supplementary Fig. 5. Structural comparison between open Kv7.2 conformations**. Superposition between our homology-based model (HM), in green, and the cryo-EM open structures: (**A)** PDB ID 8J01[19], **(B)** PDB ID 8J02[19], **(C)** PDB ID 8W4U[19], **(D)** PDB ID 8IJK[20]. RMSDs were calculated using all 115 Cα atoms of the chains retained in the simulations.

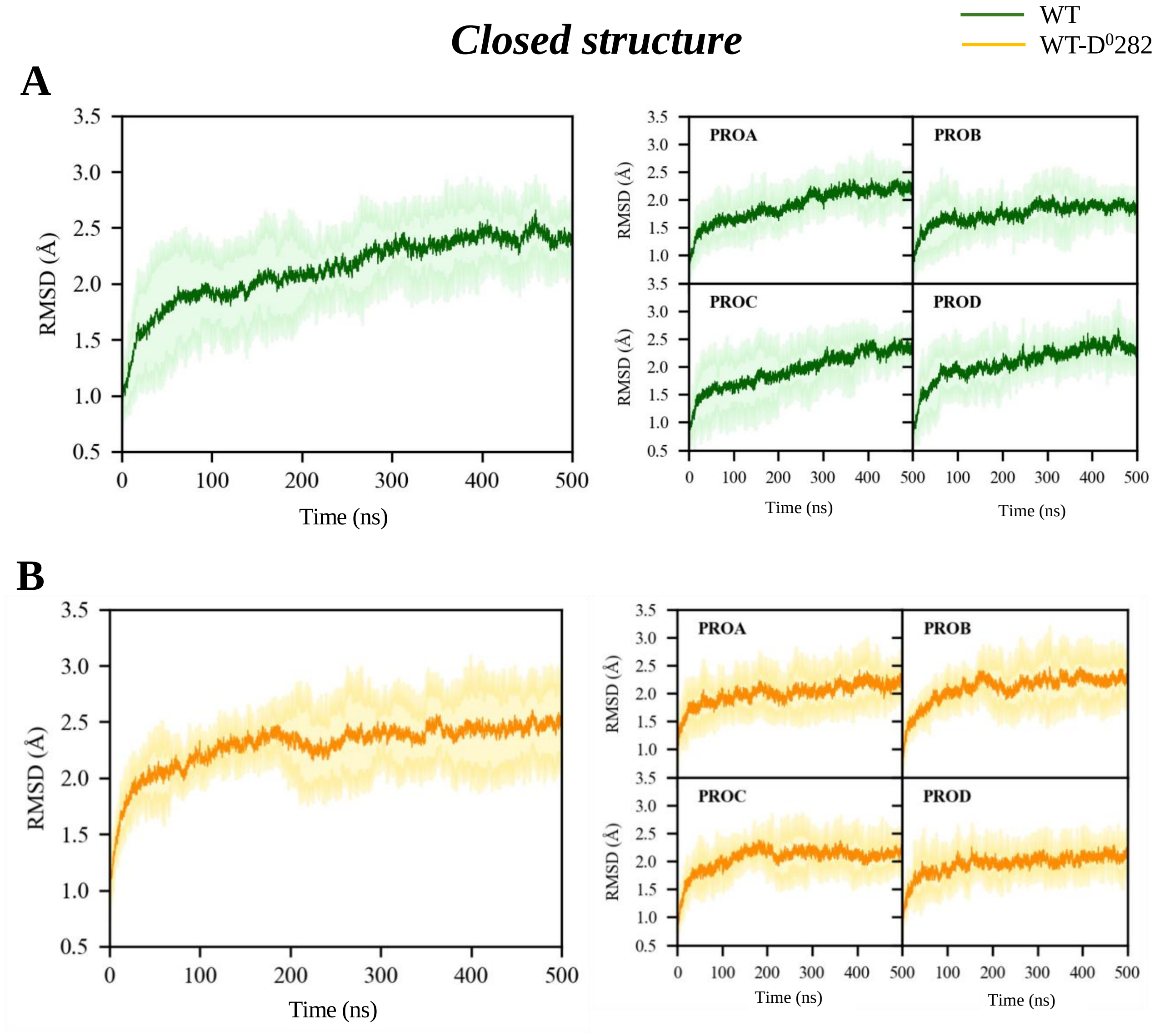


**Supplementary Fig. 6. Time evolution of TM backbone RMSD values**. RMSD values were averaged over all replicas of the Kv7.2 channels with a starting closed IG: **(A)** WT, **(B)** WT-D$^0$282. Shaded areas represent standard deviations. The plots on the right report values for each subunit, labeled as PROA, PROB, PROC, PROD.

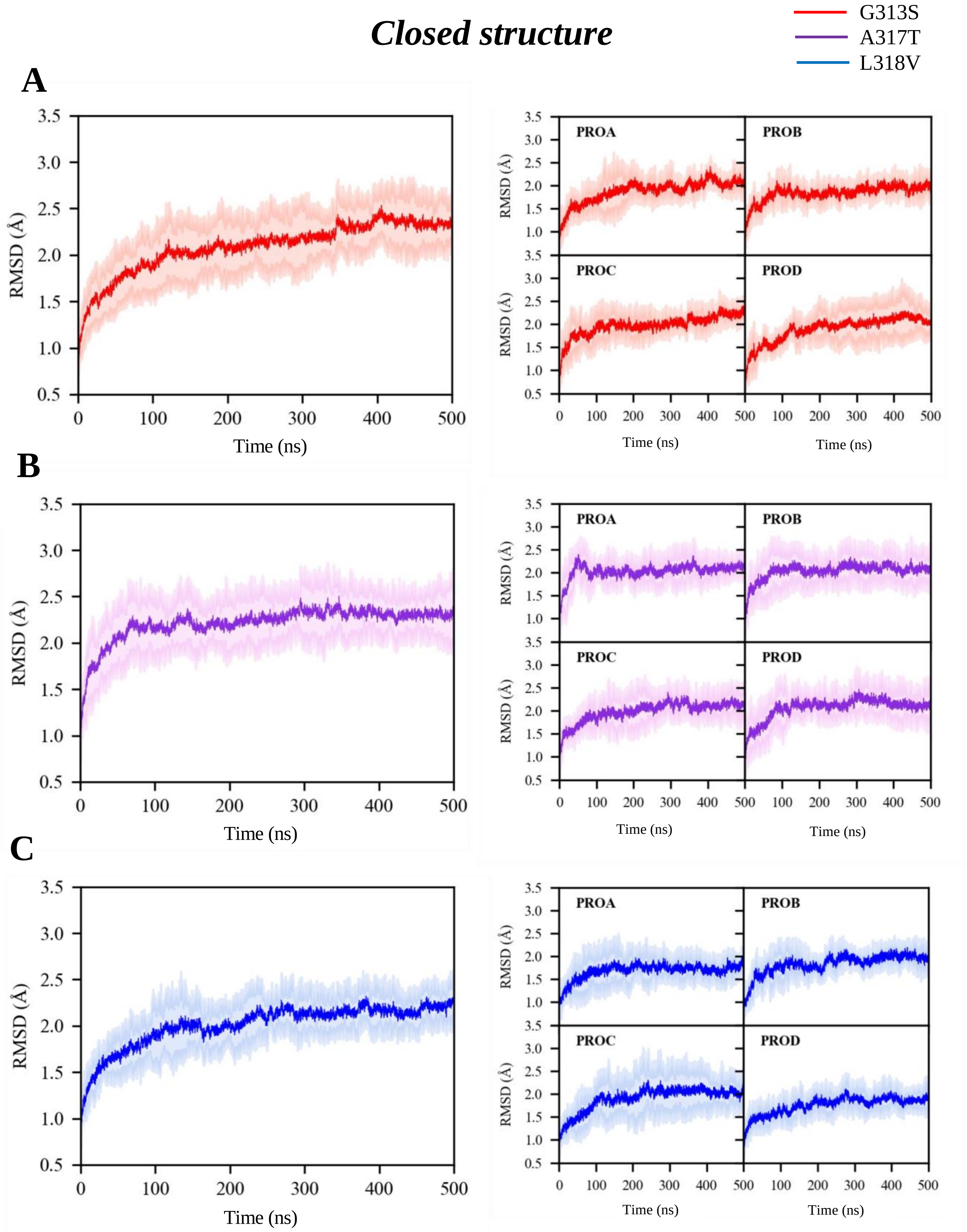


**Supplementary Fig. 7. Time evolution of TM backbone RMSD values.** RMSD values were averaged over all replicas of the Kv7.2 channels with a starting closed IG: **(A)** G313S, **(B)** A317T, **(C)** L318V. Shaded areas represent standard deviations. The plots on the right report values for each subunit, labeled as PROA, PROB, PROC, PROD.

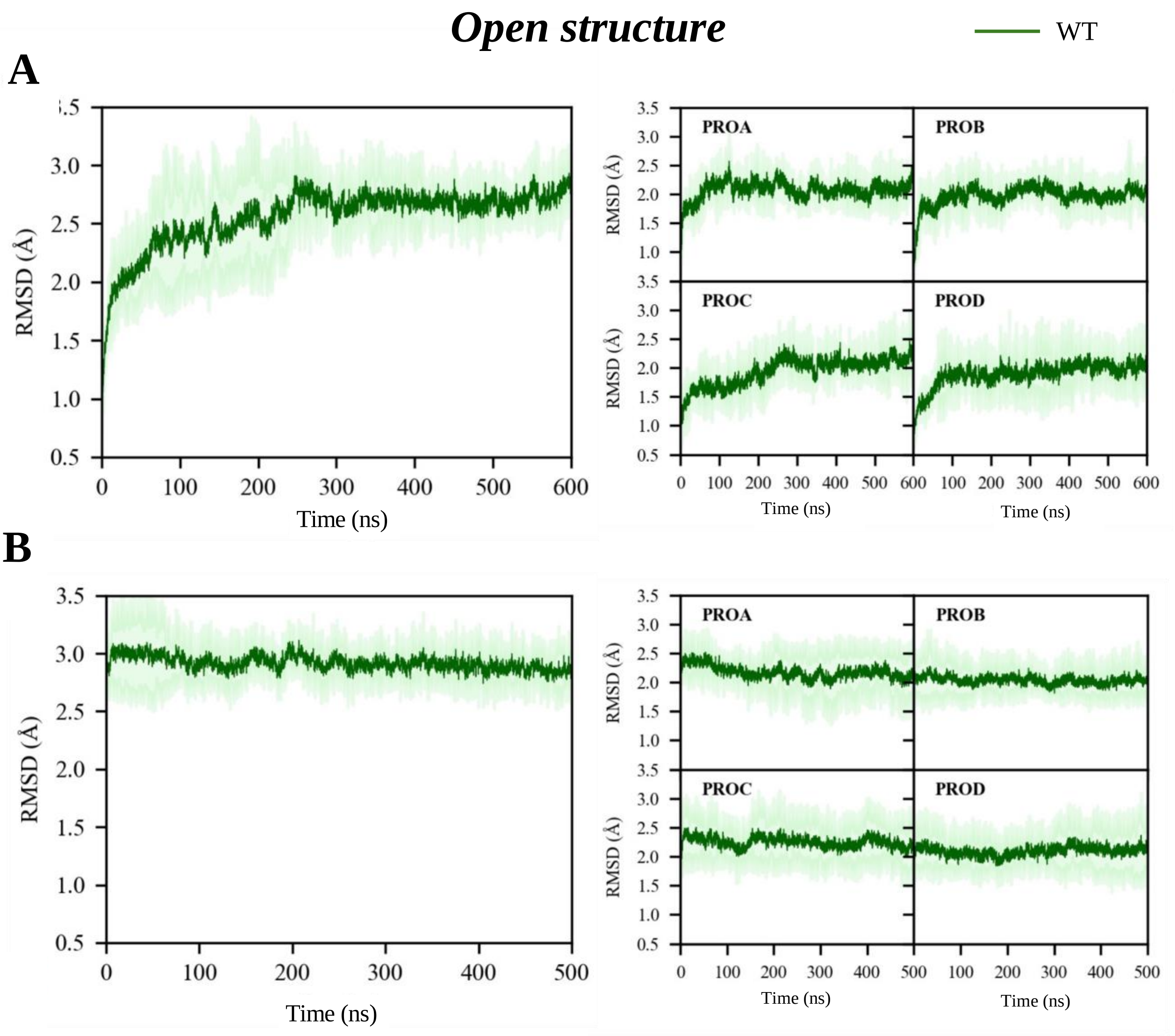


**Supplementary Fig. 8. Time evolution of TM backbone RMSD values**. RMSD values were averaged over all replicas of the WT Kv7.2 channel with an open IG: **(A)** WT restrained equilibration stage (see Materials and Methods for details), **(B)** WT production stage. Shaded areas represent standard deviations. The plots on the right report values for each subunit, labeled as PROA, PROB, PROC, PROD.

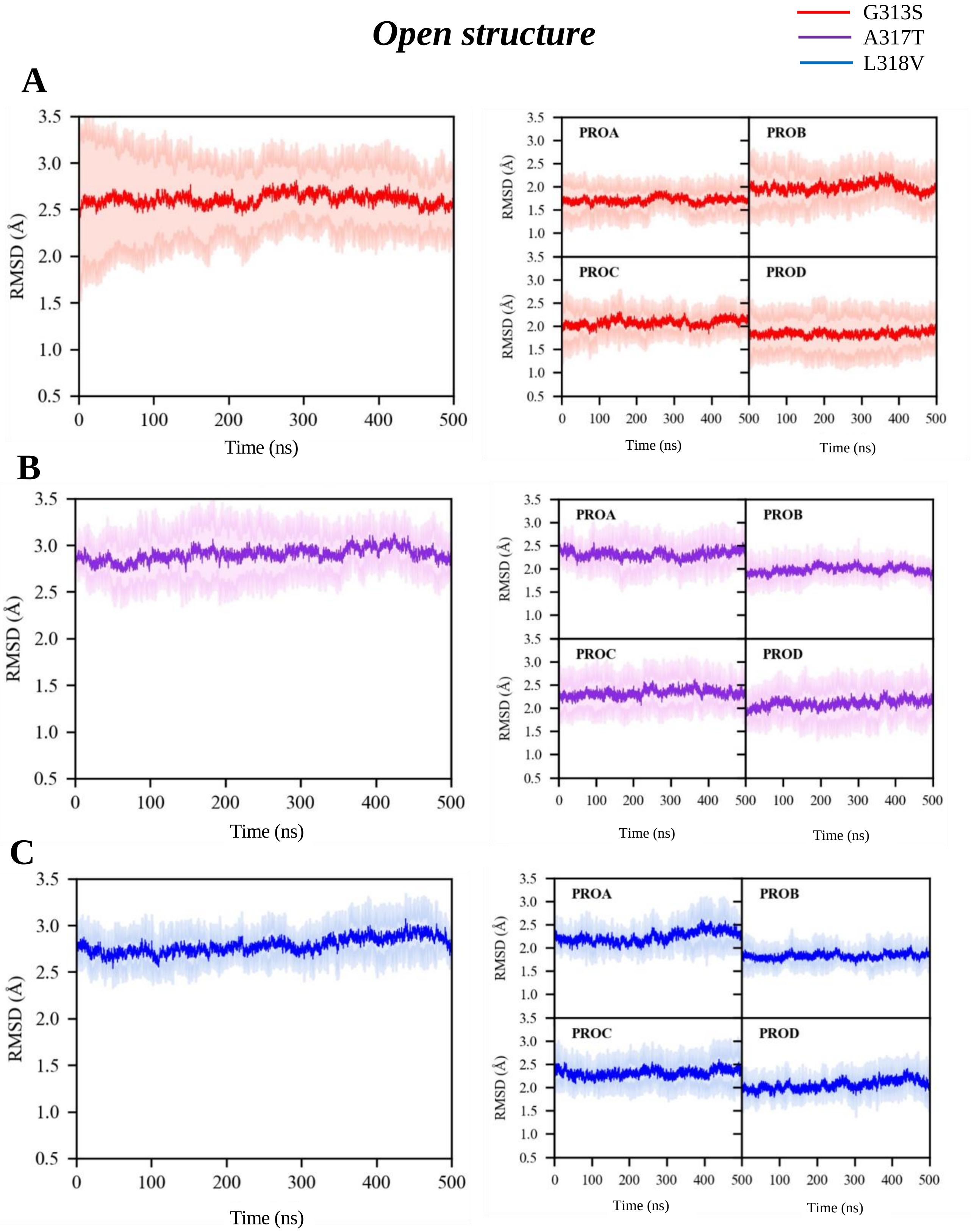


**Supplementary Fig. 9. Time evolution of TM backbone RMSD values.** RMSD values were averaged over all replicas of the Kv7.2 mutated channels with an open IG: **(A)** G313S **(B)** A317 **(C)** L318V. Shaded areas represent standard deviations. The plots on the right report values for each subunit, labeled as PROA, PROB, PROC, PROD.

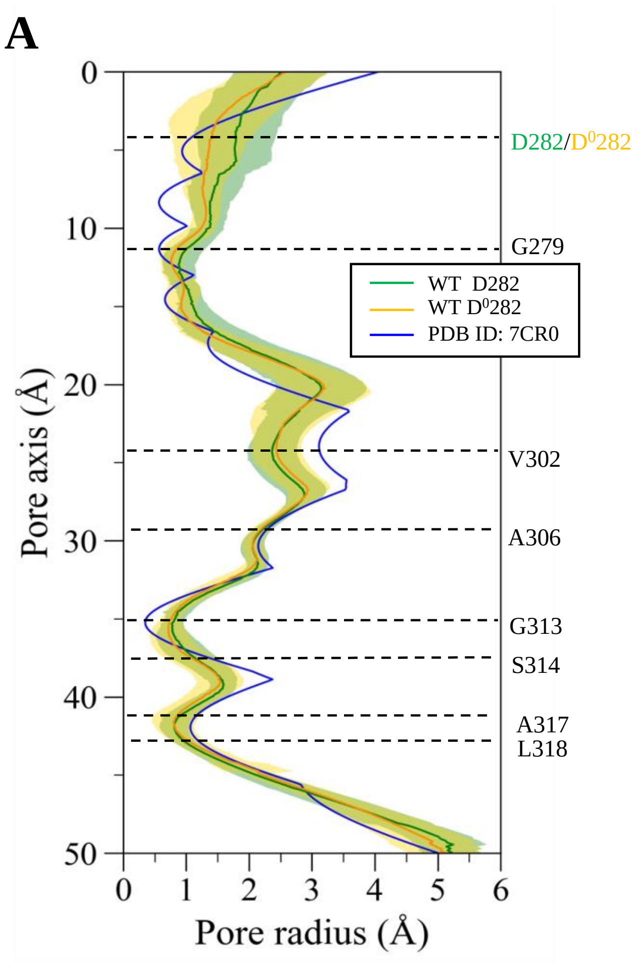


**Supplementary Fig. 10. Pore profiles of WT Kv7.2. (A)** Closed IG, radius profiles calculated from simulations with charged (green) and neutral (orange) D282. Data are averaged over the five replicated trajectories. Blue line: radius profile of the cryo-EM hKv7.2 structure (PDB ID: 7CR0). **(B)** Open IG, channel radius profile of the WT channel (green) averaged over the five replicated trajectories. Blue line: radius profile of the cryo-EM structure of the homolog hKv7.1 (PDB ID: 6V01) used as template.

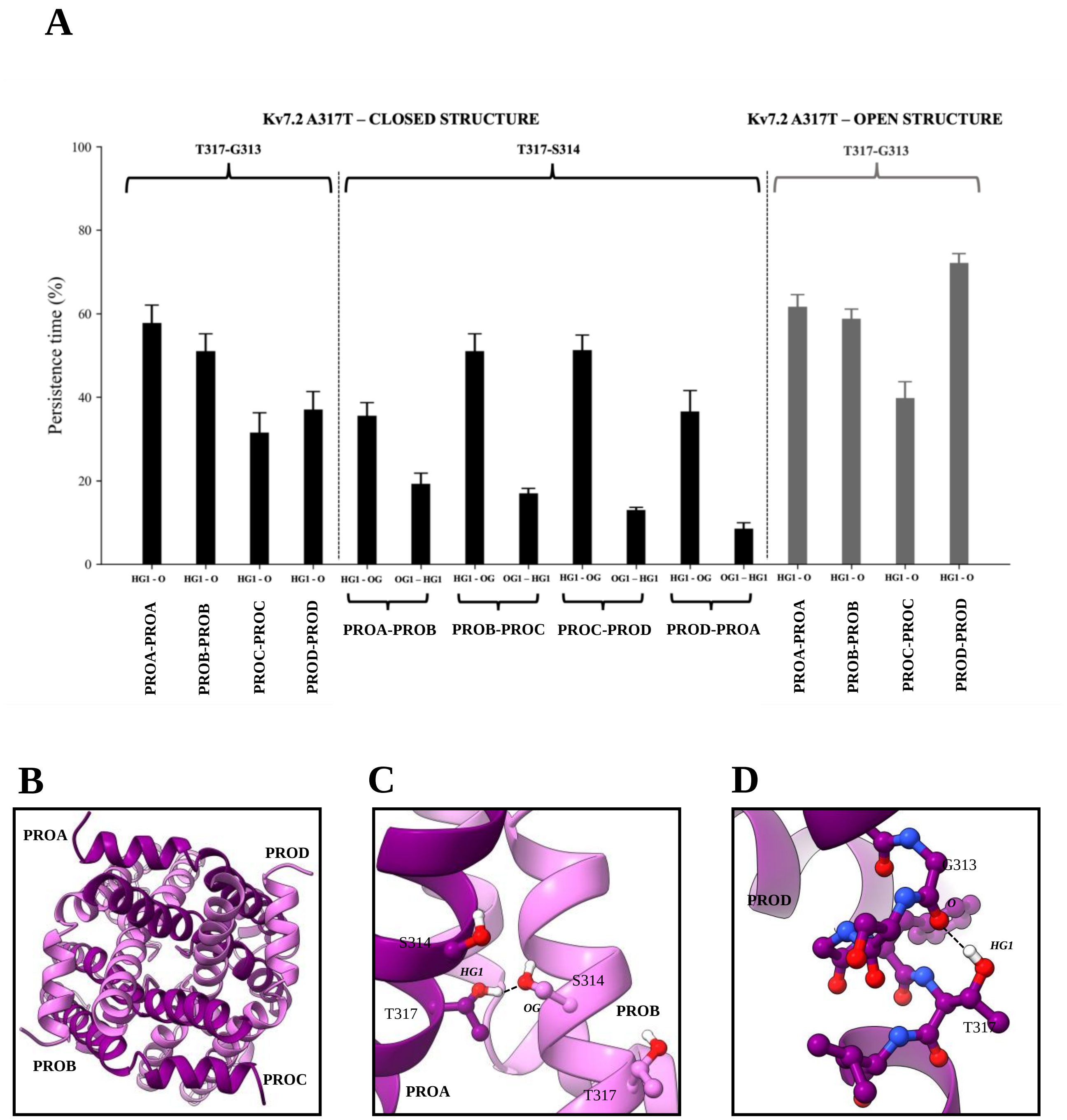


**Supplementary Fig. 11. HB analysis for the A317T simulations. (A)** Persistence of the HBs (reported as percentage of the aggregated simulation time) in the closed (black bars) and open (gray bars) structures during MD runs. The interactions are classified based on the atoms involved and the subunits they belong to. Atom names follow the PDB CHARMM notation: O, backbone amide oxygen; HG1, the hydrogen atom bound to an oxygen in γ position with respect to the backbone amide carbon atom; OG, the hydroxyl oxygen in serine (oxygen atom in γ position with respect to the backbone amide carbon atom), and OG1 the hydroxyl oxygen in threonine (again with O atom in γ position with respect to the backbone amide carbon atom). **(B)** Snapshot of the Kv7.2 pore with a closed IG, viewed from the cytosol. Equilibrated structure at $t$ = 160 ns of an MD run. **(C)** Representative structures showing the HBs formed between the T317-S314 sidechains and **(D)** T317 sidechain and G313 backbone.

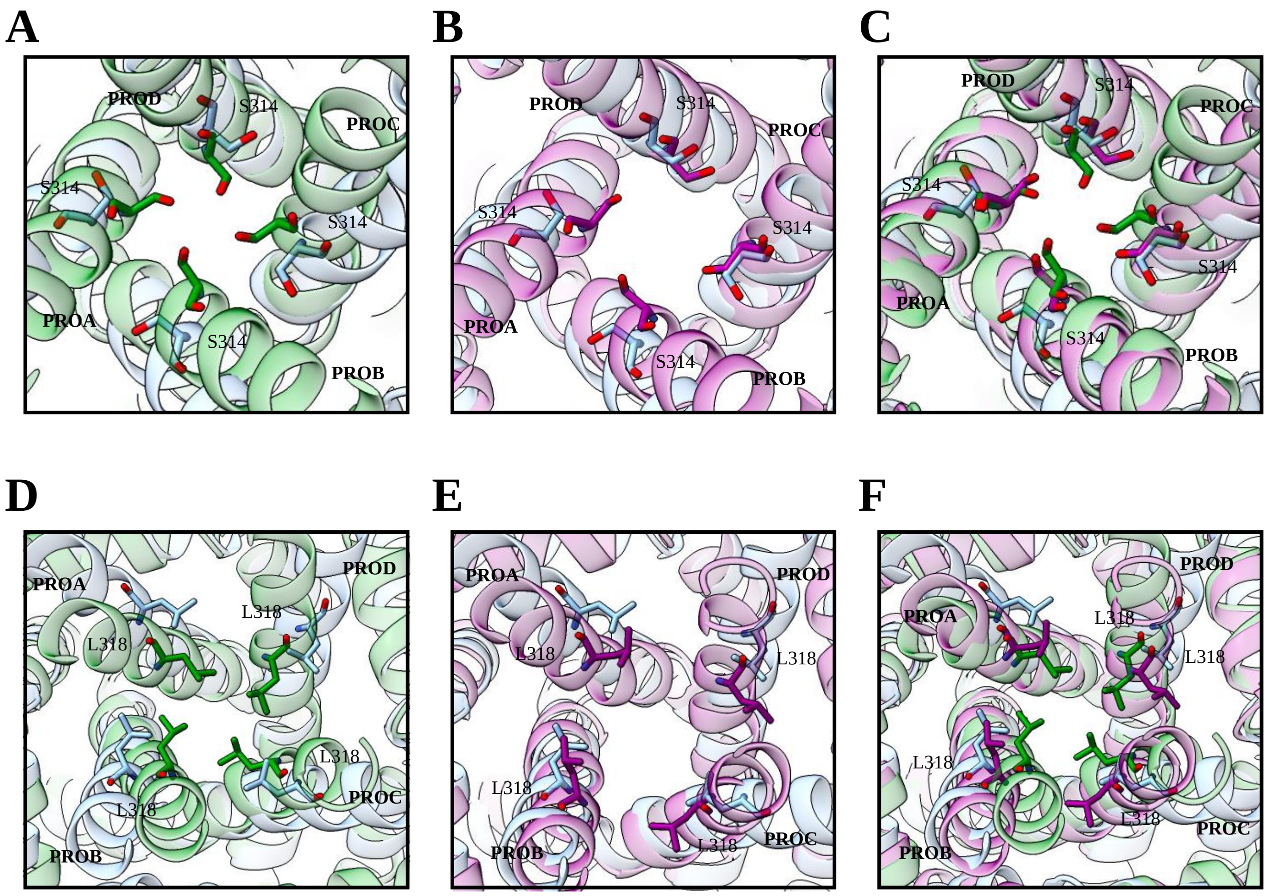


**Supplementary Fig. 12. Comparison of S314 and L318 orientation in different Kv7.2 structures. (A,B,D,E)** Green chain: WT closed IG (after 500 ns of MD); purple chain: A317T mutant, closed IG (after 500 ns); light-blue: open WT bound to ebio1 (PDB ID: 8IJK). **(C,F)** The three structures together.

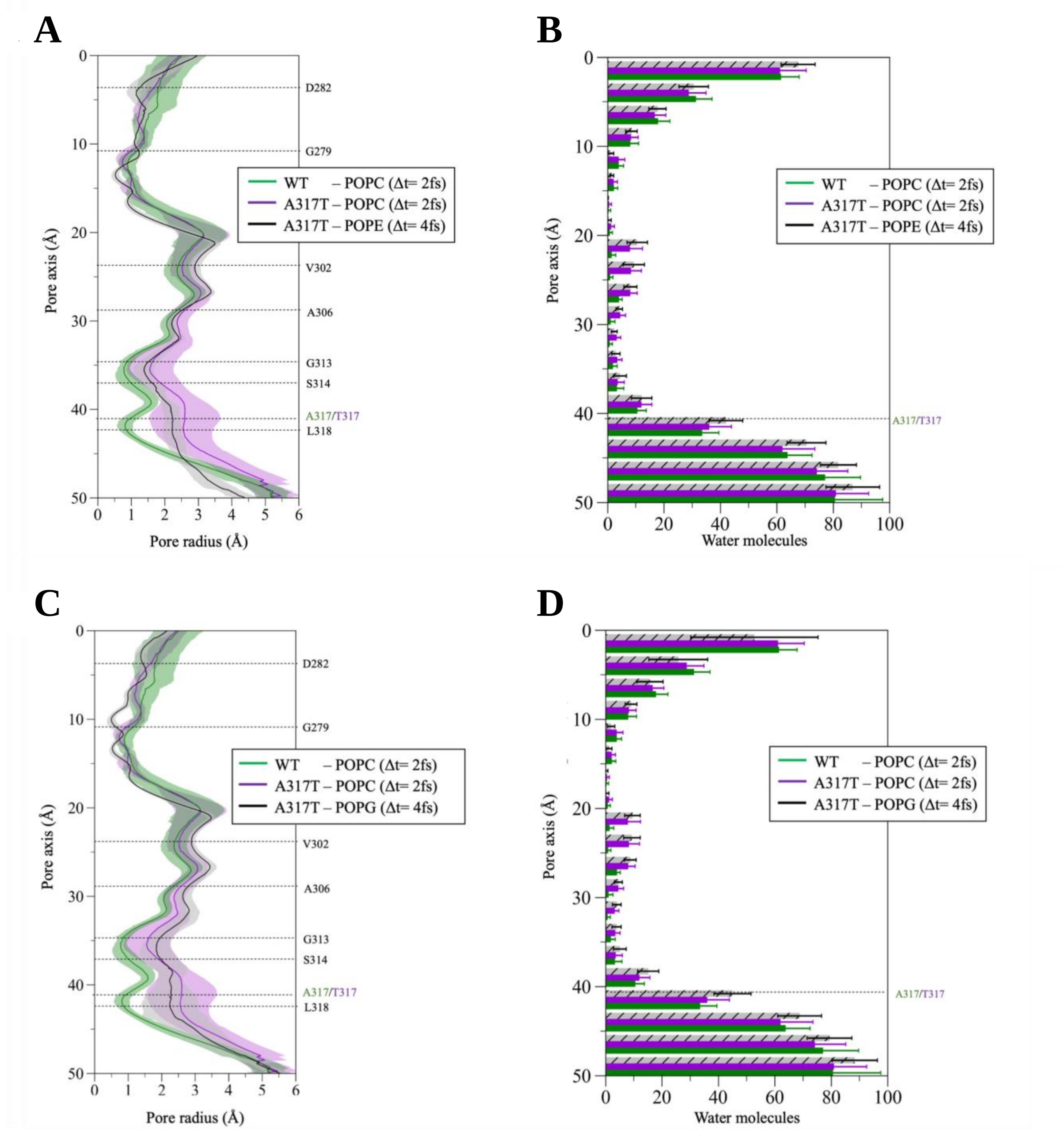


**Supplementary Fig. 13. Simulations of A317T in different membrane models. (A)** Radius profile (*left*) and histograms of water occupancy (*right*) along the axis of the A317T Kv7.2 closed channel, embedded in a POPE membrane (HMR set-up). Results are compared to those of the WT (green) and A317 in POPC channel shown in Main text. **(B)** Same as **(A)** but with the channel embedded in a POPG membrane.

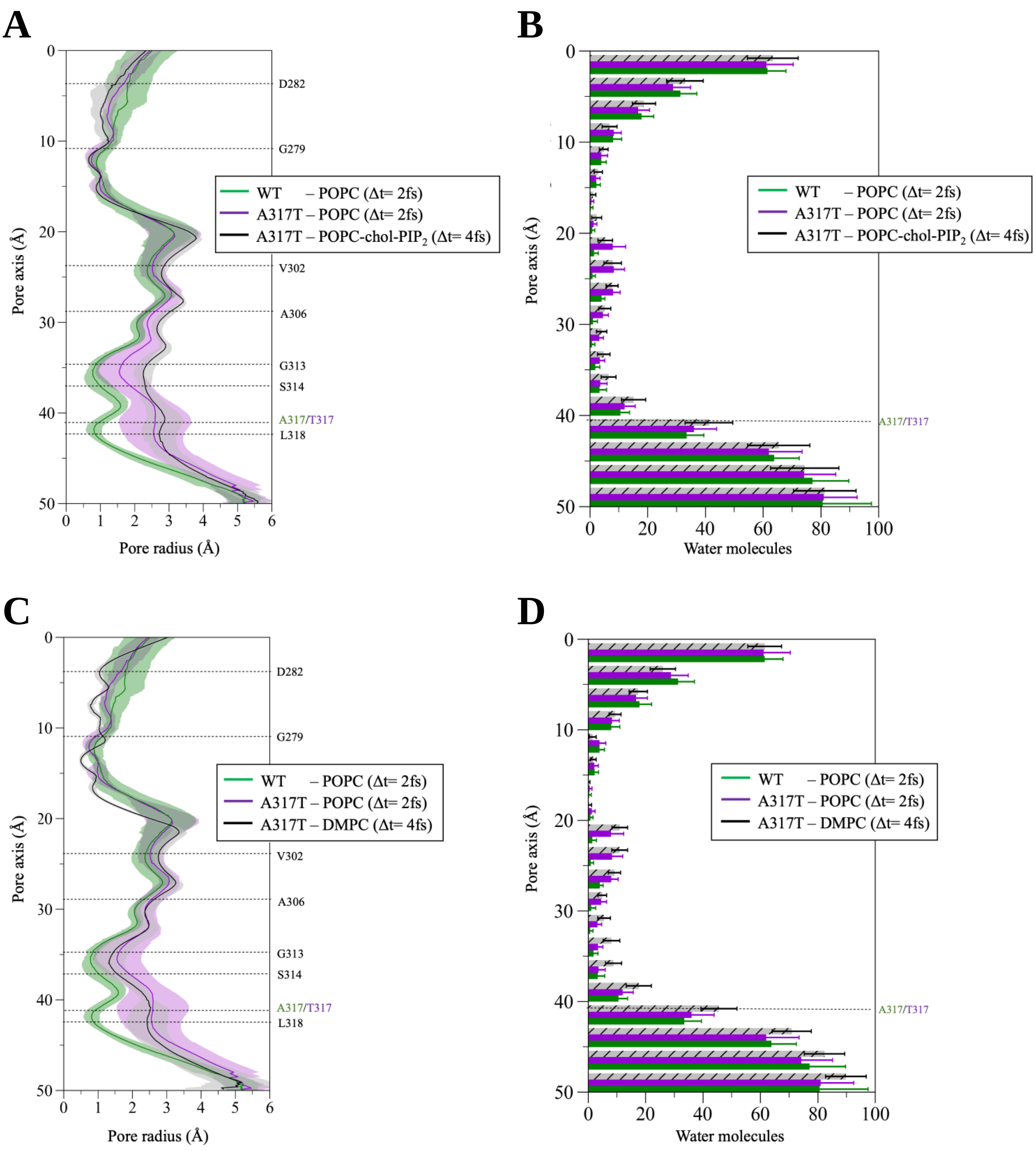


**Supplementary Fig. 14. Simulations of A317T in different membrane models. (A)** Radius profile (*left*) and histograms of water occupancy (*right*) along the axis of the A317T Kv7.2 closed channel, embedded in a POPC-chol-$PIP_2$ membrane (HMR set-up). Results are compared to those of the WT (green) and A317 in POPC channel shown in Main text. **(B)** Same as **(A)** but with the channel embedded in a DMPC membrane.

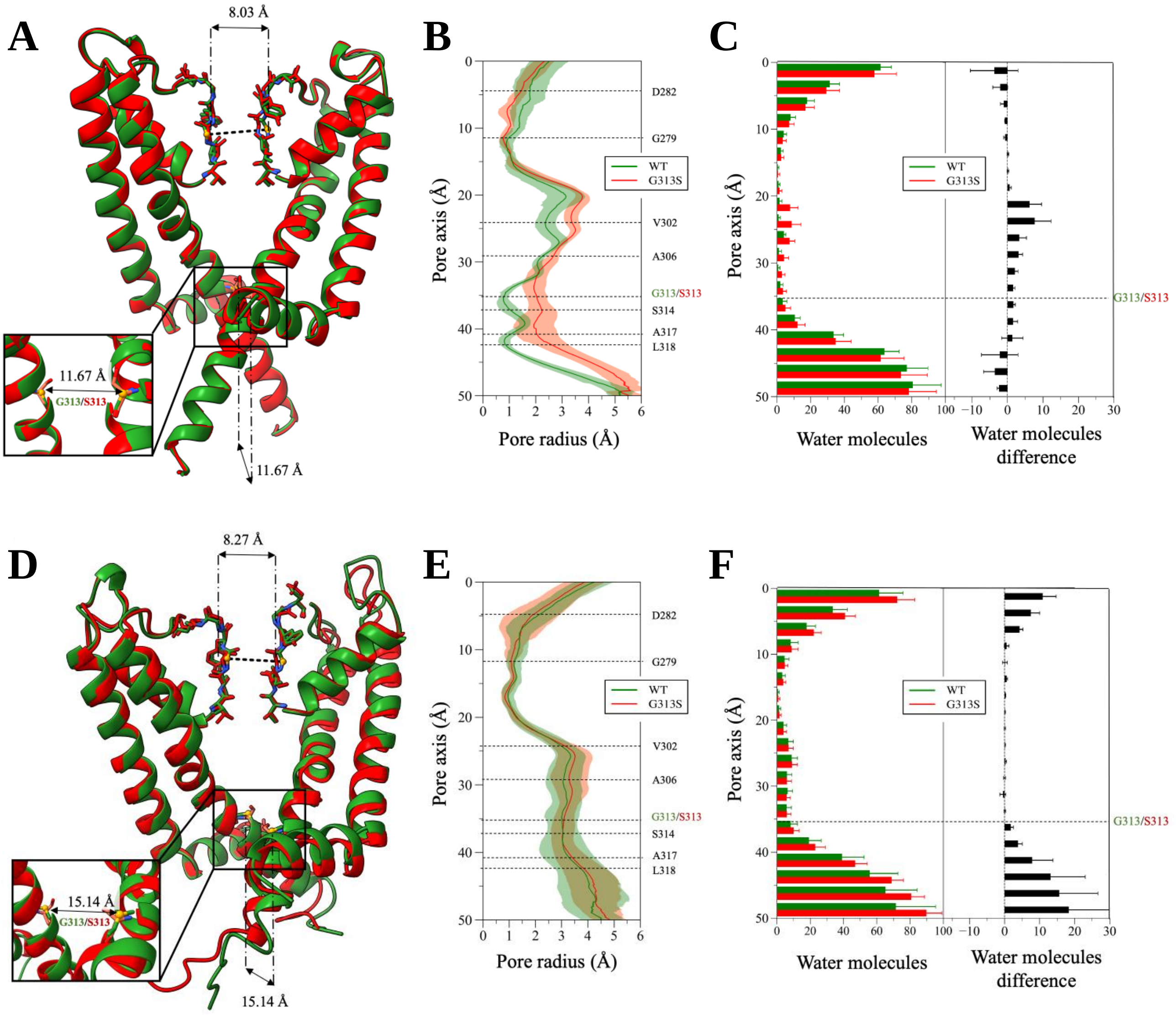


**Supplementary Fig. 15. MD simulations of the S313G Kv7.2 variant. (A)** Superposition of WT (green) and G313S (red) Kv7.2 representative closed structures after equilibration and before MD production simulations. **(B)** Channel radius profiles along the pore axis of the two proteins averaged over all simulated replicas. Shaded regions indicate standard deviations. **(C)** Distribution of water molecules along the channel axis. Average and standard deviations are calculated over all replicas. **(D-F)** Same as **(A-C)** but for the open structures.

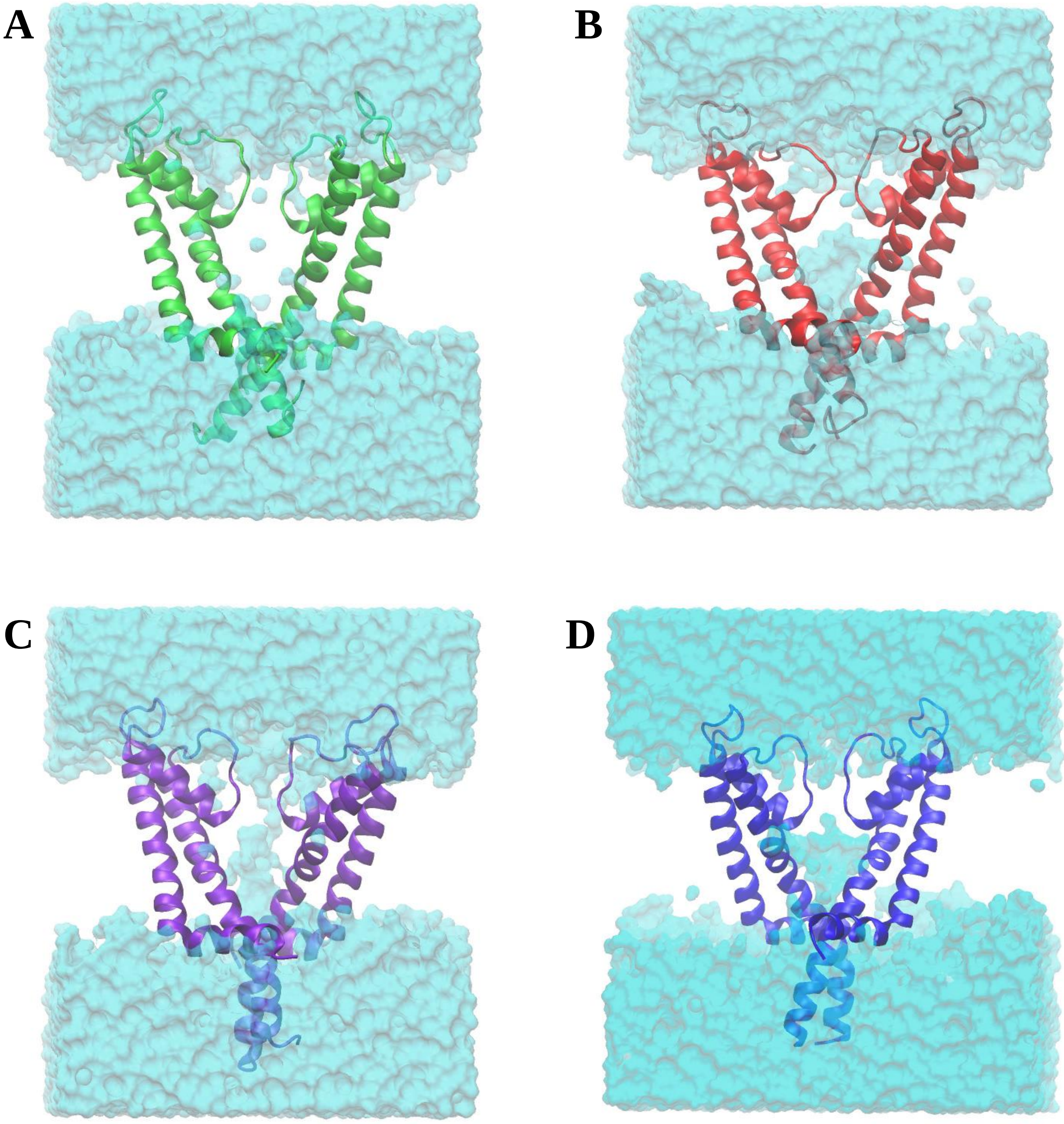


**Supplementary Fig. 16**. **Kv7.2 pore cavity hydration**. **(A)** WT, **(B)** G313S, **(C)** A317T, **(D)** L318V. For each system, we report an equilibrated structure extracted at 150 ns. Water molecules are represented as cyan surfaces. Lipid molecules and ions are not reported for clarity.

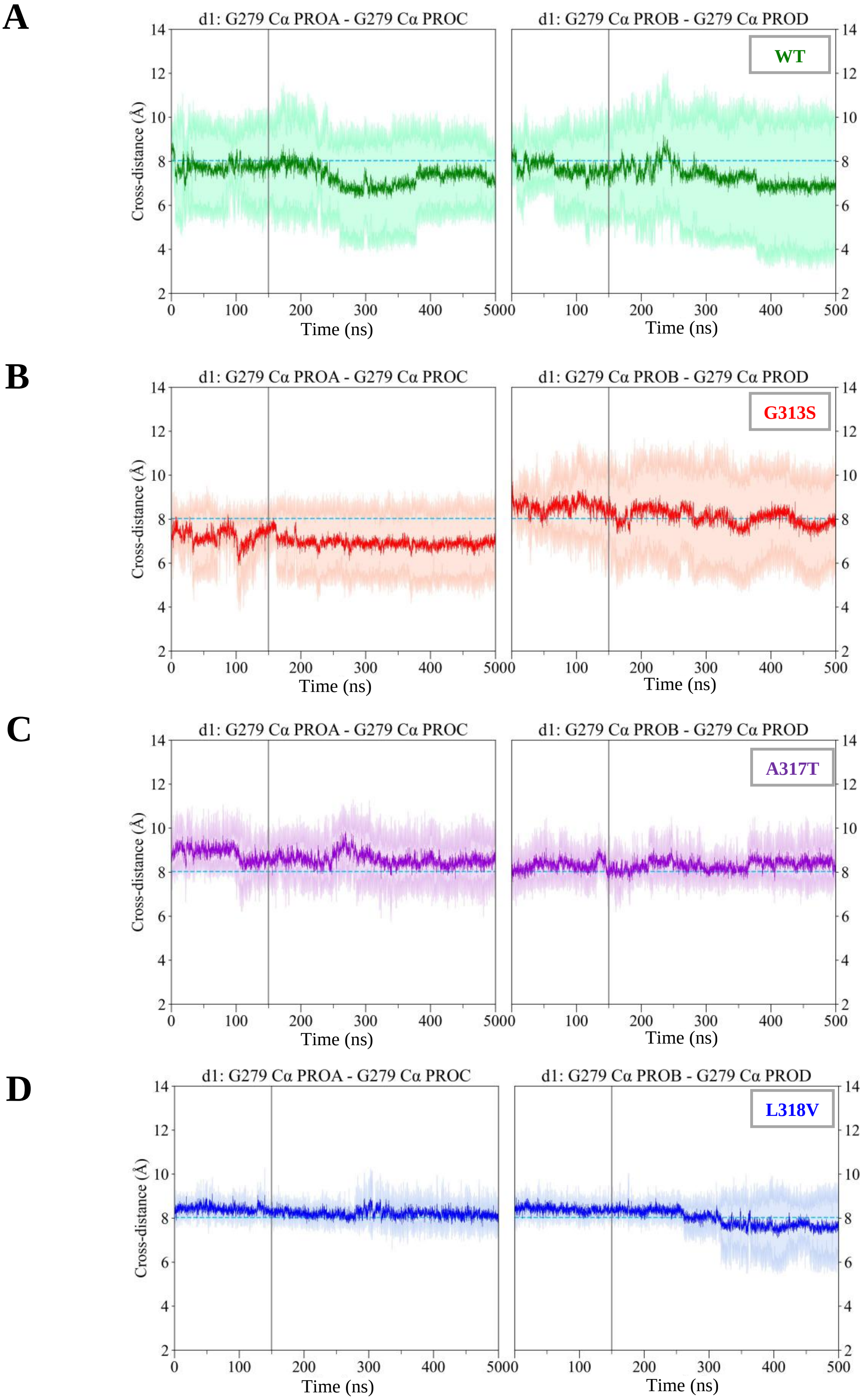


**Supplementary Fig. 17. Closed IG d1 distances.** Time evolution (from the cumulative MD simulations) of the average d1 cross distance (darker solid line) and standard deviation (lighter line) for WT **(A)**, G313S **(B)**, A317T **(C)** and L318V **(D)**. For each system, the reference value (d1=8.03 Å) measured in the starting WT structure (PDB ID: 7CR0) is reported as a dashed cyan line.

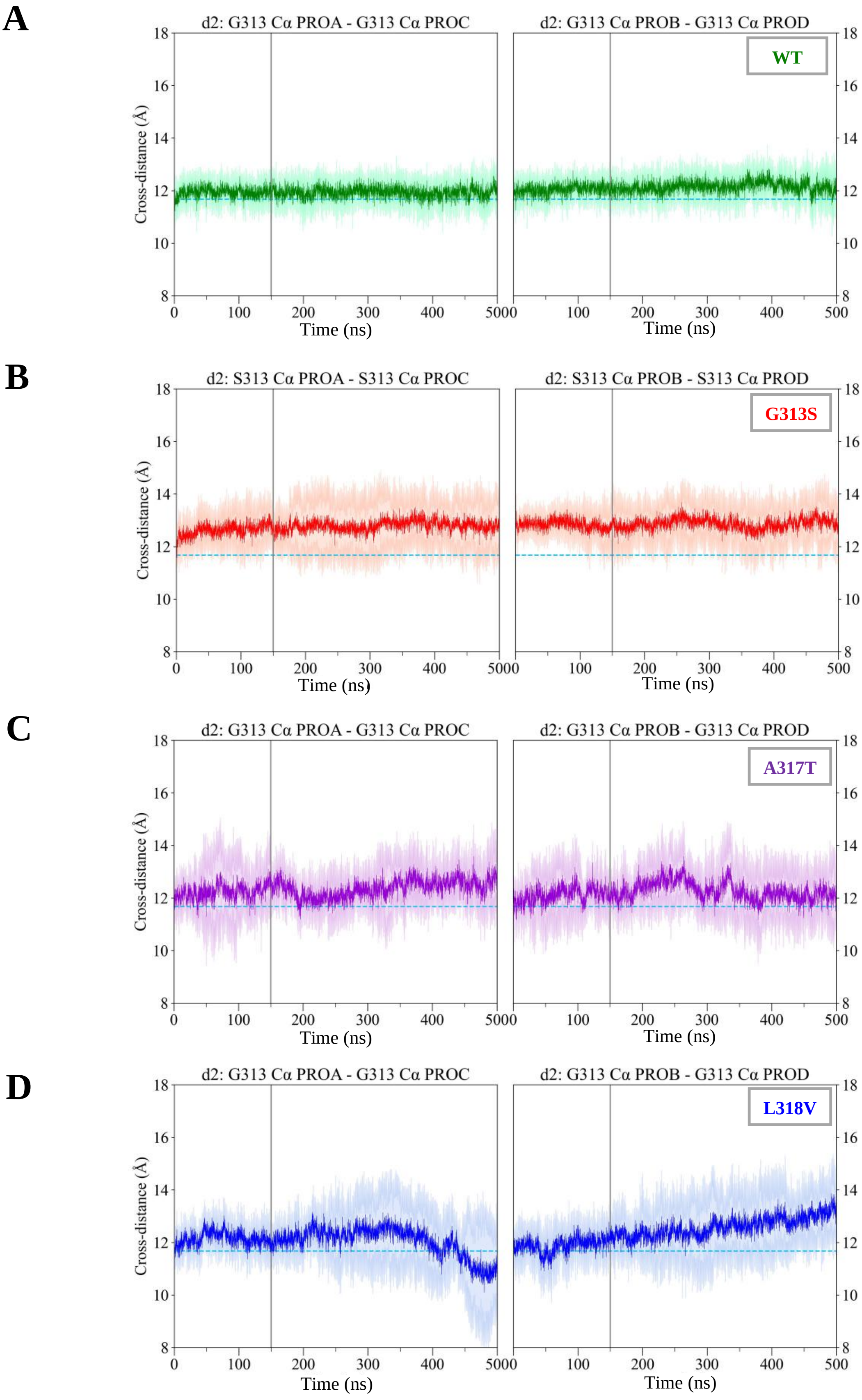


**Supplementary Fig. 18. Closed IG d2 distances.** Time evolution (from the cumulative MD simulations) of the average d2 cross distances (darker solid line) and standard deviation (shaded areas) for WT **(A)**, G313S **(B)**, A317T **(C)** and L318V **(D)**. For each system, the reference value (d2=11.67 Å) measured in the starting WT structure (PDB ID: 7CR0) is reported as a dashed cyan line.

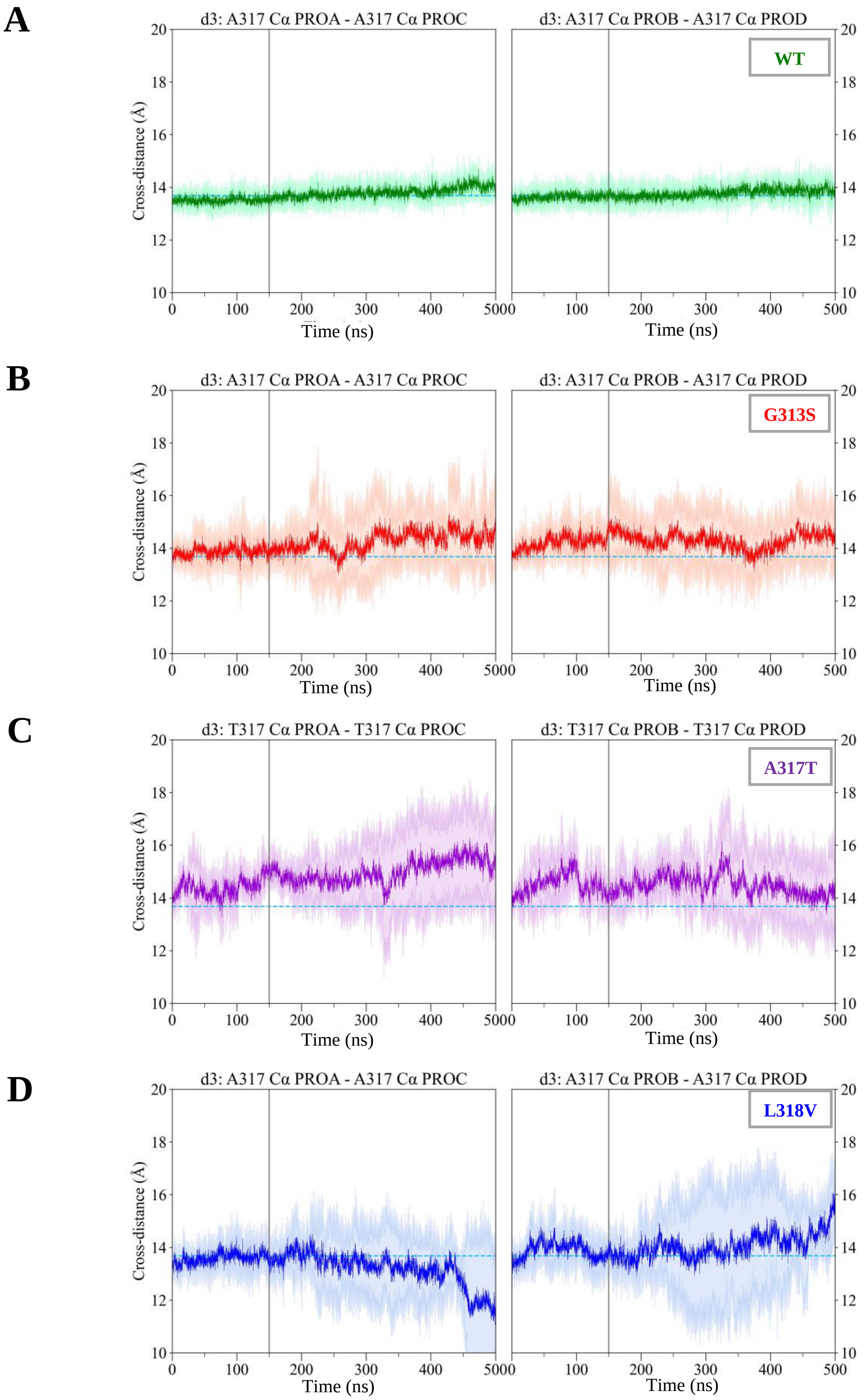


**Supplementary Fig. 19. Closed IG d3 distances.** Time evolution (from the cumulative MD simulations) of the average d3 cross distances (darker solid line) and standard deviation (shaded areas) for WT **(A)**, G313S **(B)**, A317T **(C)** and L318V **(D)**. For each system, the reference value (d3=13.69 Å) measured in the starting WT structure (PDB ID: 7CR0) is reported as a dashed cyan line.

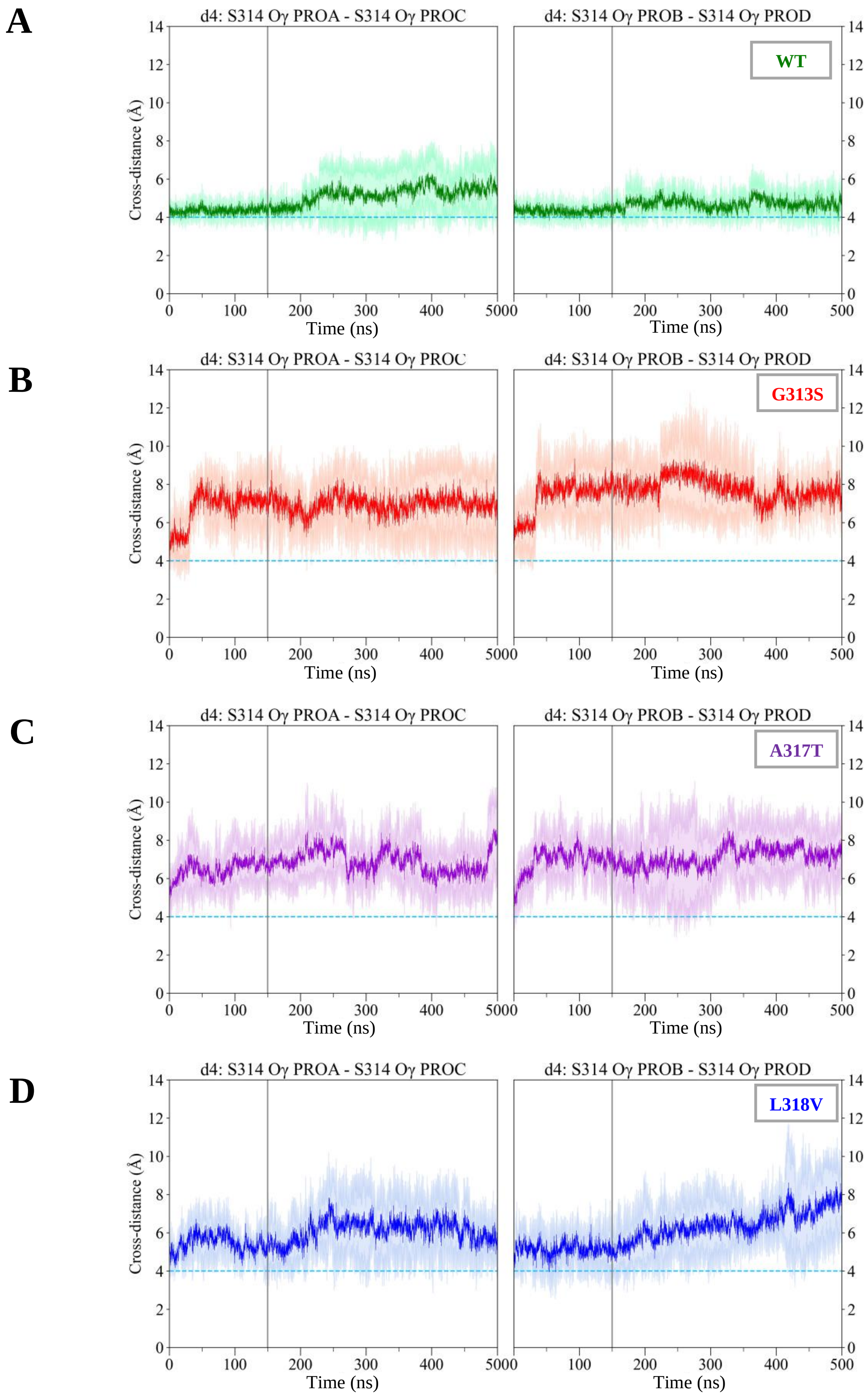


**Supplementary Fig. 20. Closed IG d4 distances.** Time evolution (from the cumulative MD simulations) of the average d4 cross distances (darker solid line) and standard deviation (shaded areas) for WT **(A)**, G313S **(B)**, A317T **(C)** and L318V **(D)**. For each system, the reference value (d4=4.00 Å) measured in the starting WT structure (PDB ID: 7CR0) is reported as a dashed cyan line.

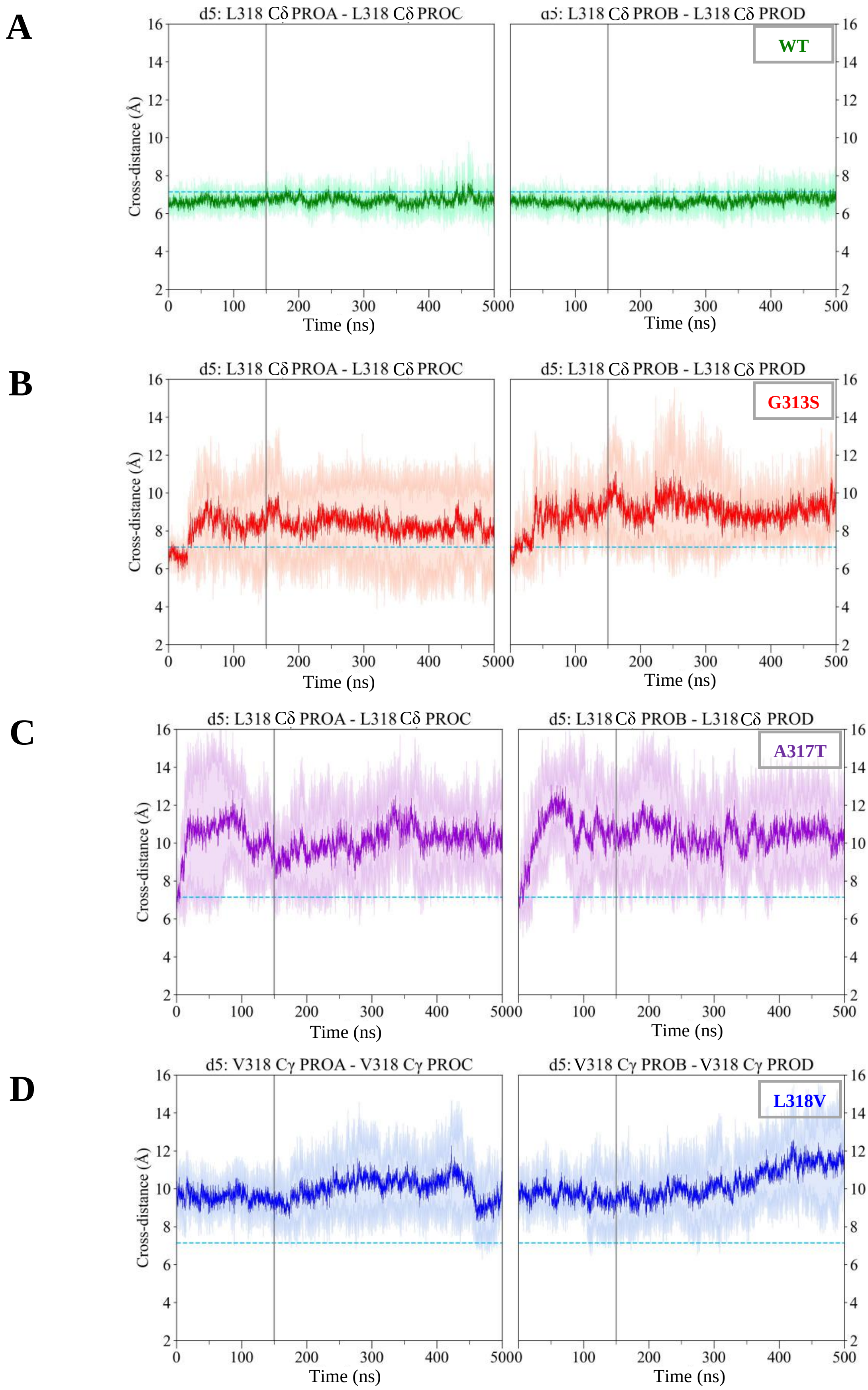


**Supplementary Fig. 21. Closed IG d5 distances.** Time evolution (from the cumulative MD simulations) of the average d5 cross distances (darker solid line) and standard deviation (shaded areas) for WT **(A)**, G313S **(B)**, A317T **(C)** and L318V **(D)**. For each system, the reference value (d5=7.14 Å) measured in the starting WT structure (PDB ID: 7CR0) is reported as a dashed cyan line.

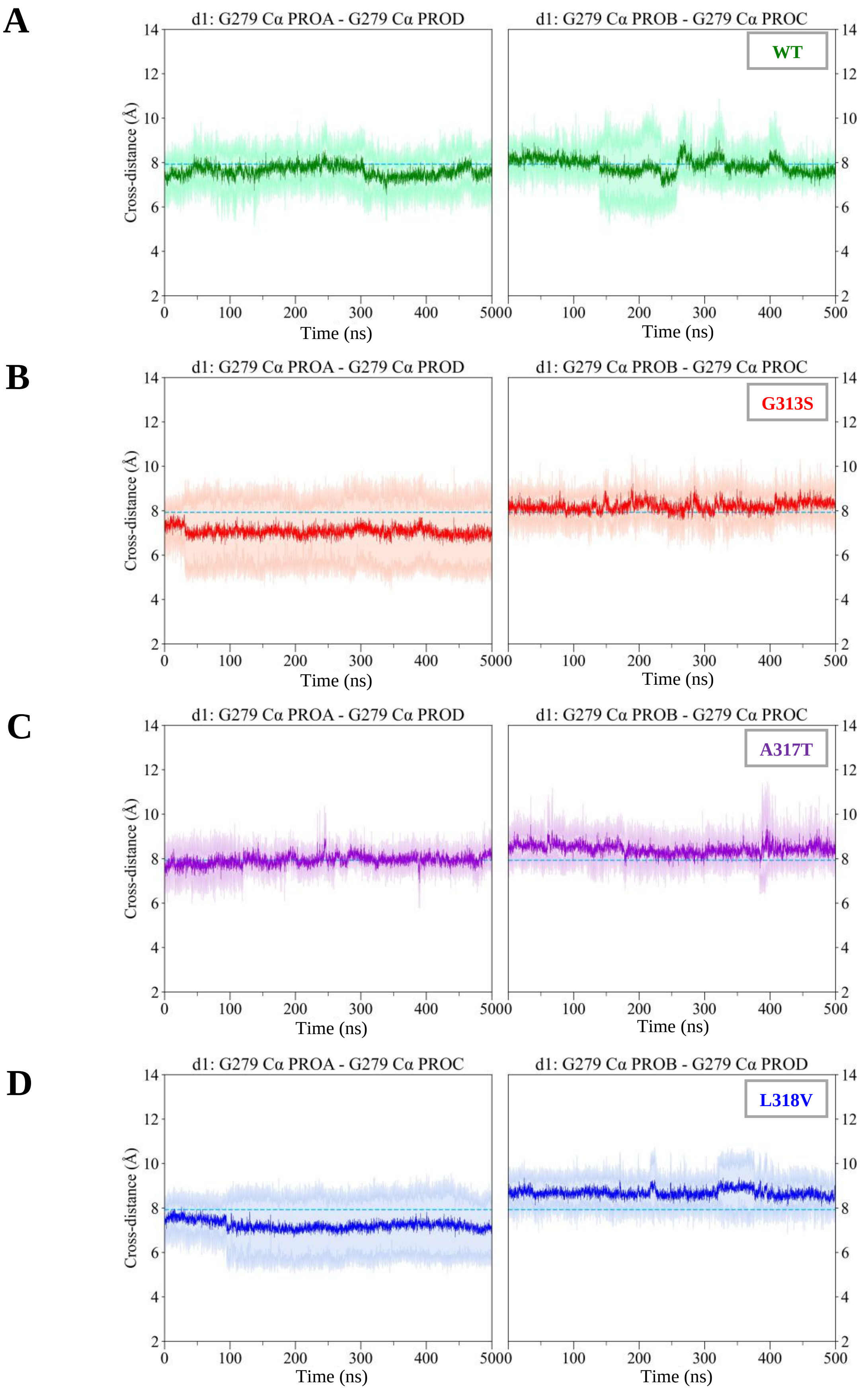


**Supplementary Fig. 22. Open IG d1 distances.** Time evolution (from the cumulative MD simulations) of the average d1 cross distances (darker solid line) and standard deviation (shaded areas) for WT **(A)**, G313S **(B)**, A317T **(C)** and L318V **(D)**. For each system, the reference value (d1=7.97 Å) measured in the homology-based model of the open Kv7.2 pore is reported as a dashed cyan line.

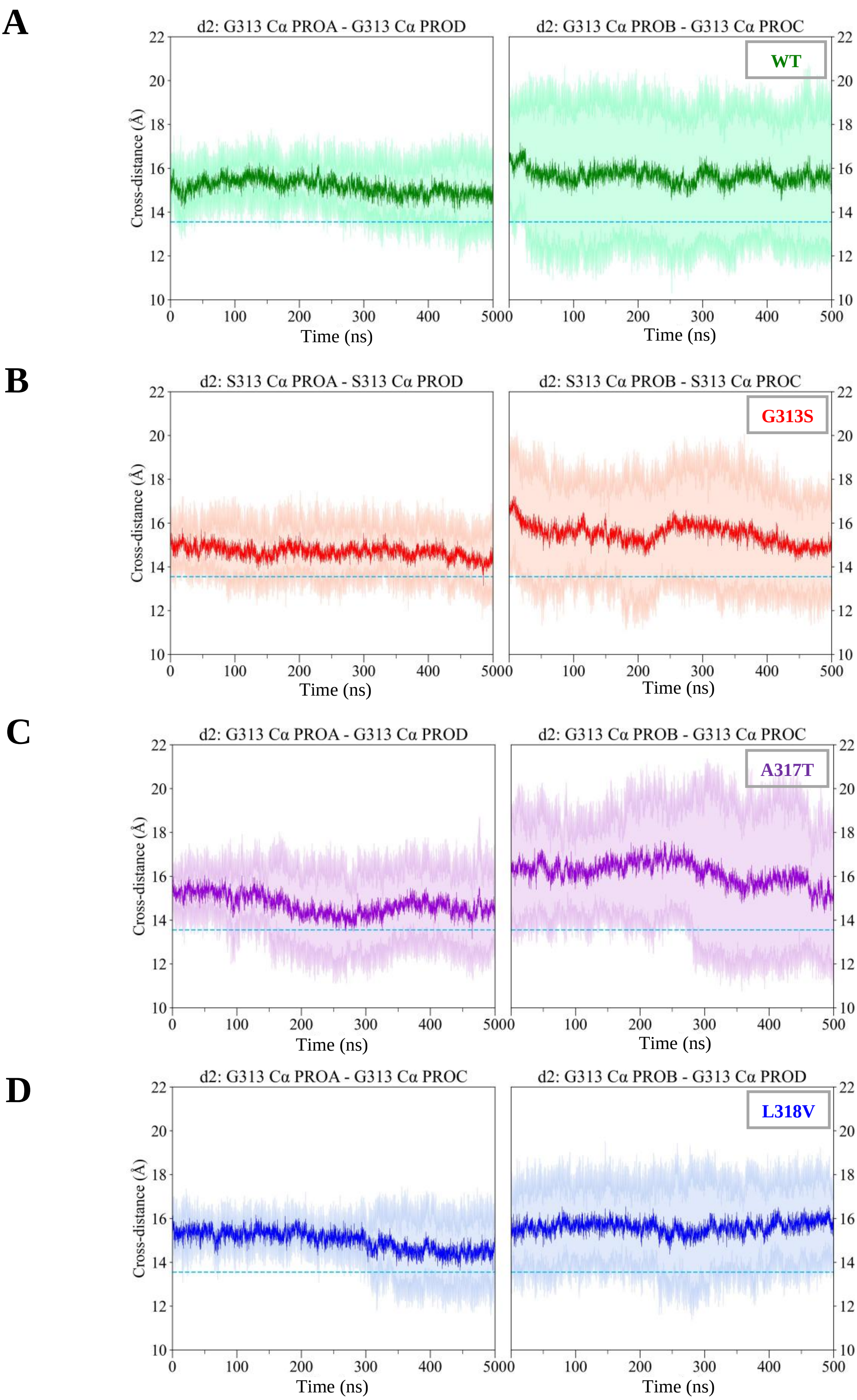


**Supplementary Fig. 23. Open IG d2 distances.** Time evolution (from the cumulative MD simulations) of the average d2 cross distances (darker solid line) and standard deviation (shaded areas) for WT **(A)**, G313S **(B)**, A317T **(C)** and L318V **(D)**. For each system, the reference value (d2 =13.55 Å) measured in the homology-based model of the Kv7.2 pore is reported as a dashed cyan line.

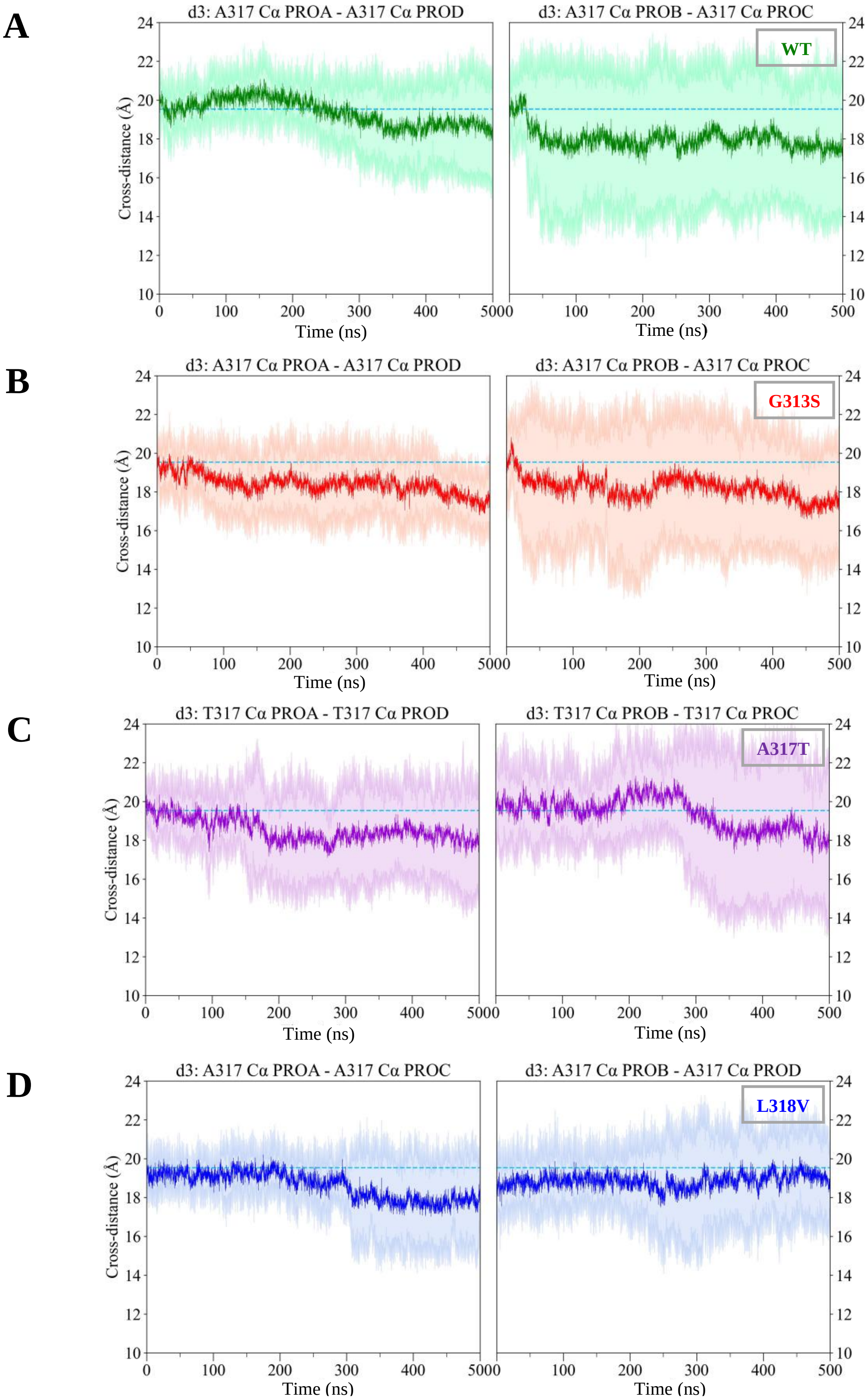


**Supplementary Fig. 24. Open IG d3 distances.** Time evolution (from the cumulative MD simulations) of the average d3 cross distances (darker solid line) and standard deviation (shaded areas) for WT **(A)**, G313S **(B)**, A317T **(C)** and L318V **(D)**. For each system, the reference value (d3=19.62 Å) measured in the homology-based model of the Kv7.2 pore is reported as a dashed cyan line. **25.** Time evolution (from the cumulative MD simulations) of the average d1 cross distances (darker solid line) and standard deviation (shaded areas) between the Cα atoms of the A317 residues (or T317 in the case of the first mutant) belonging to the two diagonally opposed monomers for WT **(A),** G313S **(B),** A317T **(C),** L318V **(D),** starting from a closed IG. For each system the reference value (d4=4.00 Å) measured in the starting WT structure (PDB ID: 7CR0) is reported as a dashed cyan line.

### BIBLIOGRAPHY


1. Jo, Sunhwan, Taehoon Kim, and Wonpil Im. Automated builder and database of protein/membrane complexes for molecular dynamics simulations. *PloS one* 2.9, e880 (2007).
2. Lomize, M. A., Pogozheva, I. D., Joo, H., Mosberg, H. I. & Lomize, A. L. OPM database and PPM web server: resources for positioning of proteins in membranes. *Nucleic Acids Research* 40, D370–D376 (2012).
3. Phillips, J. C. *et al.* Scalable molecular dynamics with NAMD. *J Comput Chem* 26, 1781–1802 (2005).
4. Best, R. B. *et al.* Optimization of the additive CHARMM all-atom protein force field targeting improved sampling of the backbone $\phi$, $\psi$ and side-chain $\chi$1 and $\chi$2 dihedral angles. *J. Chem. Theory Comput.* 8, 3257–3273 (2012).
5. Huang, J. & MacKerell, A. D. CHARMM36 all-atom additive protein force field: validation based on comparison to NMR data. *J Comput Chem* 34, 2135–2145 (2013).
6. Klauda, J. B. *et al.* Update of the CHARMM all-atom additive force field for lipids: validation on six lipid types. *J. Phys. Chem. B* 114, 7830–7843 (2010).
7. Jorgensen, W. L., Chandrasekhar, J., Madura, J. D., Impey, R. W. & Klein, M. L. Comparison of simple potential functions for simulating liquid water. *The Journal of Chemical Physics* 79, 926–935 (1983).
8. Noskov, S. Y. & Roux, B. Control of ion selectivity in LeuT: two Na+ binding sites with two different mechanisms. *Journal of Molecular Biology* 377, 804–818 (2008).
9. Luo, Y. & Roux, B. Simulation of osmotic pressure in concentrated aqueous salt solutions. *J. Phys. Chem. Lett.* 1, 183–189 (2010).
10. Venable, R. M., Luo, Y., Gawrisch, K., Roux, B. & Pastor, R. W. Simulations of anionic lipid membranes: development of interaction-specific ion parameters and validation using NMR Data. *J. Phys. Chem. B* 117, 10183–10192 (2013).
11. Darden, T., York, D. & Pedersen, L. Particle mesh Ewald: An N.log(N) method for Ewald sums in large systems. *The Journal of Chemical Physics* 98, 10089–10092 (1993).
12. Steinbach, P. J. & Brooks, B. R. New spherical-cutoff methods for long-range forces in macromolecular simulation. *Journal of Computational Chemistry* 15, 667–683 (1994).
13. Ryckaert, J.-P., Ciccotti, G. & Berendsen, H. J. C. Numerical integration of the cartesian equations of motion of a system with constraints: molecular dynamics of n-alkanes. *Journal of Computational Physics* 23, 327–341 (1977)
14. Miyamoto, S. & Kollman, P. A. Settle: An analytical version of the SHAKE and RATTLE algorithm for rigid water models. *Journal of Computational Chemistry* 13, 952–962 (1992).
15. Feller, S. E., Zhang, Y., Pastor, R. W. & Brooks, B. R. Constant pressure molecular dynamics simulation: The Langevin piston method. *The Journal of Chemical Physics* 103, 4613–4621 (1995).
16. Martyna, G. J., Tobias, D. J. & Klein, M. L. Constant pressure molecular dynamics algorithms. *The Journal of Chemical Physics* 101, 4177–4189 (1994).
17. Huang, Jing, et al. CHARMM36m: an improved force field for folded and intrinsically disordered proteins. *Nature methods* 14.1, 71-73 (2017).
18. Balusek, C. *et al.* Accelerating membrane simulations with hydrogen mass repartitioning. *J Chem Theory Comput* 15, 4673–4686 (2019).
19. Ma, D. *et al.* Ligand activation mechanisms of human KCNQ2 channel. *Nat Commun* 14, 6632 (2023).
20. Zhang, S. *et al.* A small-molecule activation mechanism that directly opens the KCNQ2 channel. *Nat Chem Biol* 1–10 (2024)
